# Discovery and characterization of *LNCSOX17* as an essential regulator in human endoderm formation

**DOI:** 10.1101/2022.09.12.507139

**Authors:** Alexandro Landshammer, Adriano Bolondi, Helene Kretzmer, Christian Much, René Buschow, Alina Rose, Hua-Jun Wu, Sebastian Mackowiak, Bjoern Braendl, Pay Giesselmann, Rosaria Tornisiello, Krishna Mohan Parsi, Jack Huey, Thorsten Mielke, David Meierhofer, René Maehr, Denes Hnisz, Franziska Michor, John L. Rinn, Alexander Meissner

## Abstract

Long non-coding RNAs (lncRNAs) have emerged as fundamental regulators in various biological processes, including embryonic development and cellular differentiation. Despite much progress over the past decade, the genome-wide annotation of lncRNAs remains incomplete and many known non-coding loci are still poorly characterized. Here, we report the discovery of a previously not annotated lncRNA that is transcribed upstream of the *SOX17* gene and located within the same topologically associating domain. We termed it *LNCSOX17* and show that it is induced following SOX17 activation but its expression is more tightly restricted to early definitive endoderm. Loss of *LNCSOX17* affects crucial functions independent of SOX17 and leads to an aberrant endodermal transcriptome, signaling pathway deregulation and epithelial to mesenchymal transition defects. Consequently, cells lacking the lncRNA cannot further differentiate into more mature endodermal cell types. We identified and characterized *LNCSOX17* as an essential new actor in early human endoderm, thereby further expanding the list of functionally important non-coding regulators.

## INTRODUCTION

To date, nearly 28.000 long non-coding RNAs (lncRNAs) have been reported in the human genome, but less than 1% (∼150) has been functionally characterized (*1–4*). Several of those have been shown to influence cellular physiology in developmental, adult and disease contexts (*5–10*). Depending on their genomic location, lncRNAs can be classified into genic lncRNAs (overlapping with a protein-coding gene) or intergenic lncRNAs (lincRNAs; no overlap with a protein-coding gene) (*1*). Together with transcription factors and epigenetic regulators (*11–13*), lncRNAs participate in complex gene-regulatory networks by fine-tuning gene expression in a precise and controlled manner (*14*). In particular, lncRNAs have been shown to modulate gene expression at multiple levels, including chromatin structure and folding (*15*), activating neighboring (*16*) and distal (*17*) genes, affecting RNA splicing (*18*), or influencing nuclear compartmentalization (*19–21*).

More specifically, non-coding RNAs have also been shown to fine-tune the activation and function of developmental regulators, including transcription factors responsible for maintenance of pluripotency (*22–24*), mesoderm specification (*25*) and neuronal differentiation (*26*). Recent studies have also attributed critical roles for lncRNAs in the early stages of human development, in particular during definitive endoderm specification through *cis-*regulatory activity on nearby genes (*27, 28*). For instance, *LNC00261* facilitates the activation of the proximal *FOXA2* gene via association with SMAD2/3 (*27*). A mechanistically similar *cis-*regulation of *GATA6* has been shown for lncRNA *GATA6-AS1* (*28*), while the lncRNA *DIGIT* has been reported to control *GSC* in *trans-*, via the formation of BRD3-dependent phase-separated condensates (*29, 30*).

The majority of lncRNAs exhibit highly tissue-specific expression that is often more restricted than observed for protein-coding genes (*31*). Signaling molecules, including TGF-β, WNT and the JUN/JNK/AP-pathway represent critical cascades necessary for endoderm formation, inducing the expression of endodermal factors such as SOX17, GATA6 and C-X-C chemokine receptor 4 (CXCR4) (*32–34*). SOX17 is a member of the SOX-F group of transcription factors and its expression is necessary for the specification of definitive endoderm *in vitro* (*35*) and *in vivo* (*36*). Despite being an essential and well-studied factor, much remains to be understood about the regulatory elements and nuclear organization of the *SOX17* locus. In this study, we identified and characterized *LNCSOX17*, a previously not annotated lncRNA that is located 230 kb upstream of the *SOX17* gene and predominantly active in early definitive endoderm. We show that SOX17 directly binds the *LNCSOX17* promoter, but *LNCSOX17* in turn does not appear to be involved in regulating *SOX17* transcription as its depletion has no detectable impact on *SOX17* expression. Importantly, loss of *LNCSOX17* expression leads to a failure in proper definitive endoderm progression. Together with our additional characterization, we demonstrate that *LNCSOX17* is an important regulator in human definitive endoderm differentiation.

## RESULTS

### Discovery of an unannotated non-coding transcript within the *SOX17* topological domain

So far, *SOX17* is the only annotated gene located within the 336 kb *SOX17* loop-domain insulated by strong CTCF-boundaries (Figure 1A, top). However, upon closer inspection of multiple epigenetic modifications in pluripotent stem cells (hESCs and hiPSCs) and early definitive endoderm we observed a potential unannotated gene locus. In particular, the combination of histone H3 lysine 4 trimethylation (H3K4me3) and histone H3 lysine 36 trimethylation (H3K36me3) in ESC derived endoderm suggested an RNA Polymerase II driven transcript (*37, 38*). Further supporting this, matched RNA sequencing data showed a 22 kb long transcribed region approximately 230 kb upstream of *SOX17* (Figure 1A, bottom). These results combined with a strong UCSC PhyloCSF sequence conservation points to an intergenic lncRNA (lincRNA) that we subsequently termed *LNCSOX17* (Figure 1A,B). The presence of this distal *SOX17* transcript was also observed across a number of vertebrates based on stage- and tissue-matched embryonic data (Figure supplement 1A, left).

**Figure 1:**
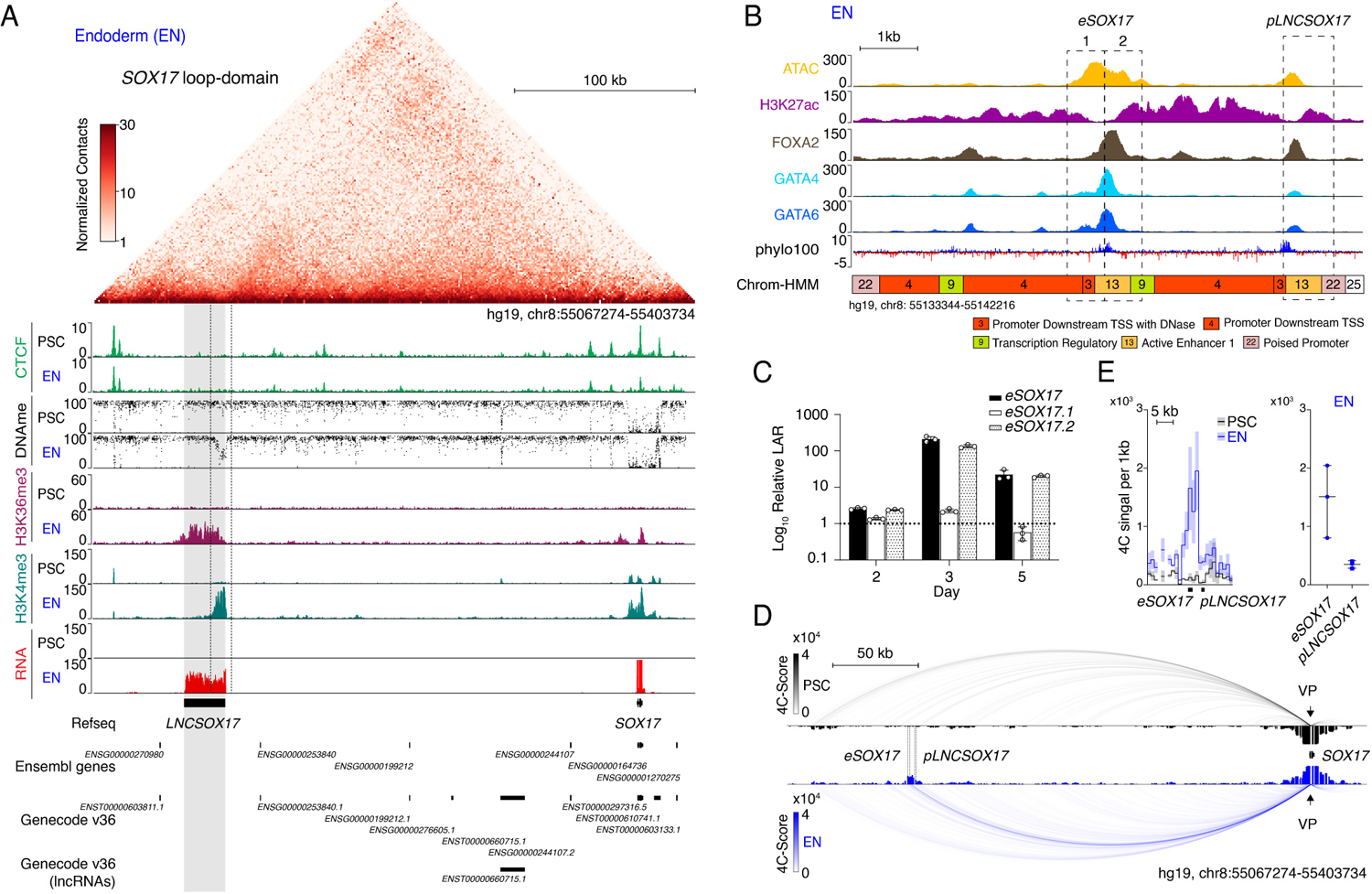
Identification of *LNCSOX17* at the human *SOX17* locus. **(A)** Normalized capture Hi-C (cHi-C) contact map of the human *SOX17* locus in endoderm cells (top panel) and chromatin immunoprecipitation sequencing (ChIP-seq) tracks of CTCF, H3K36me3 and H3K4me3 as well as whole genome bisulfite sequencing (WGBS) (Table 1) and RNA-seq profiles in PSCs and EN (bottom panel). *LNCSOX17* locus is highlighted in grey. **(B)** Zoomed in view of the *SOX17* distal regulatory element in EN cells comprising Assay for Transposase-Accessible Chromatin with high-throughput sequencing (ATAC-seq) profile and H3K27ac, FOXA2, GATA4 and GATA6 ChIP-seq (Table 1) profiles. Chrom-HMM(*136, 137*) 25-state profile is shown below the phylo100(*138, 139*) UCSC conservation track. Dashed lines indicate the two distinct regulatory elements, characterized by enriched transcription factors occupancy (*eSOX17* and *pLNCSOX17*). **(C)** Firefly luciferase assay from either *eSOX17.1* (hg19, chr8:55136923-55137557), *eSOX17.2* (hg19, chr8:55137558-55138192) or both together at day 2, 3 or 5 of EN differentiation. Values are calculated as luciferase activity ratio (LAR) between firefly and renilla signal, finally normalized on the empty vector background signal. Bars indicate mean values, error bars show standard deviation (SD) across three independent experiments. Individual data points are displayed. Raw measurements are reported in Table 1. **(D)** 4Cseq of PSC (black) and EN (blue) at the *SOX17*-locus. Normalized interaction-scores displayed as arcs and histogram-profiles utilizing the *SOX17* promoter as viewpoint (VP). **(E)** 4Cseq interactions as a zoomed in view at the *SOX17* regulatory element and corresponding quantification. In the zoomed in tracks, the line represents the median and the shaded areas depict 95% CI; in the quantification, the central line represents the median and error bars show SD across three independent experiments.

To begin exploring the locus in more detail we first set out to disentangle it from the overlapping distal regulatory element that appears to be a putative distal *SOX17* enhancer (*39*). We identified two distinct sites of transcription factor (TF) binding within a region of open chromatin specifically in definitive endoderm (Figure 1B). Although both sites show enriched UCSC PhyloCSF sequence conservation, they are also characterized by a distinguishable promoter and enhancer signature (ChromHMM state 22 and ChromHMM state 13, respectively) (Figure 1B) (*40, 41*).

The putative promoter region of *LNCSOX17* (*pLNCSOX17*) was experimentally evaluated in a luciferase assay and found to have endoderm specific activity (Figure supplement 1B). Similarly, the putative enhancer, which was further separated into two parts based on its TF occupancy profile (*eSOX17.1* and *eSOX17.2*), was also experimentally tested in a luciferase assay (Figure 1 B,C). The entire region but also *eSOX17.2* alone showed strong enhancer activity during endoderm differentiation (Figure 1C). *eSOX17’s* regulatory identity was further tested by combined epigenetic and TF binding profiling along the *LNCSOX17* locus (Figure supplement 1C).

Next, we further evaluated *eSOX17.2* using a loss of function approach. We used flanking single guide RNAs (sgRNAs) with Cas9 to generate *eSOX17.2* homozygous deletions and assessed the effect during directed endoderm differentiation (Figure supplement 1D,E). We observe a delayed activation of SOX17 and overall reduced expression of the transmembrane C-X-C chemokine receptor 4 (CXCR4), which points to a regulatory defect in the mutant cells (Figure supplement 1F,G). To further explore the physical interactions at the locus, we performed Circularized Chromosome Conformation Capture sequencing (4C-seq) on pluripotent cells and early endoderm (Figure 1D,E - Figure supplement 1H). Interestingly, we observe an enriched interaction between the *SOX17* promoter and its distal enhancer (*eSOX17*) in an endoderm specific fashion as compared to *LNCSOX17* promoter (*pLNCSOX17*) (Figure 1D,E). Therefore, we can conclude that the topologically isolated domain of *SOX17* also encompasses a distal, transcribed region driven by a promoter in close proximity but otherwise independent from a functional enhancer that interacts with the *SOX17* gene.

### *LNCSOX17* has an RNA based function during early definitive endoderm

We next investigated the expression of the non-coding transcript during endoderm differentiation with time-resolved qRT-PCR and found *LNCSOX17* expression follows *SOX17* kinetics but with an approximate 24-hour delay (Figure 2A). To explore the co-occurrence of *SOX17* and *LNCSOX17*, we compared their expression across a wide range of cell and tissue types (*n* = 44) (Figure 2B). *LNCSOX17* expression appears tightly restricted to early human definitive endoderm and uncoupled from the much broader expression of *SOX17* in many other endoderm-derived tissues (*42, 43*) (Figure 2B - Figure supplement 2A-C). Moreover, we utilized RNA-seq data from the three pluripotent stem cell derived germ layers to show that *LNCSOX17* is not expressed during mesoderm and ectoderm formation (Figure supplement 2D). scRNAseq data in the early human gastrulating embryo (*44*) further confirms *LNCSOX17’s* extreme tissue specificity *in vivo* (Figure supplement 2E).

**Figure 2:**
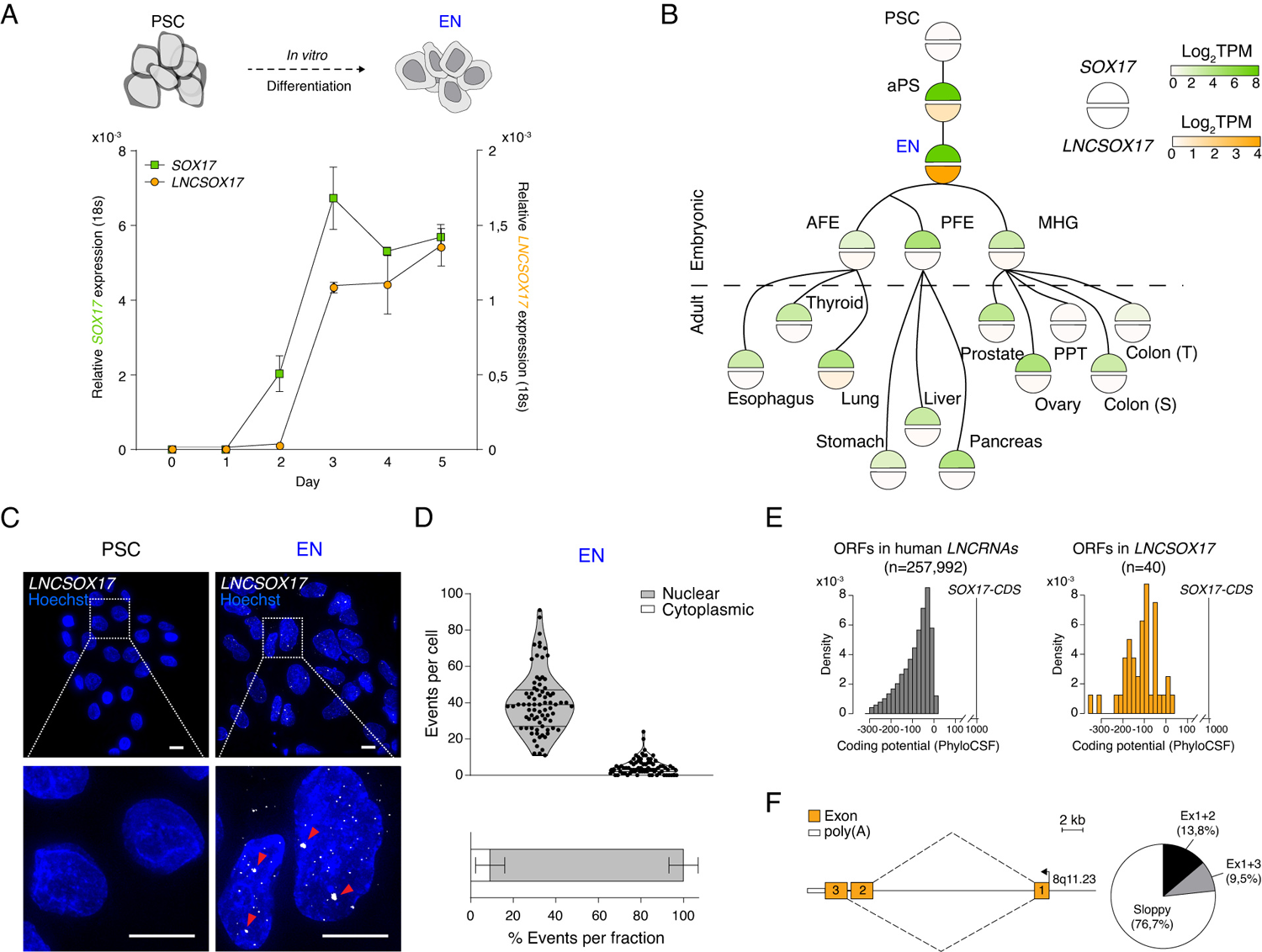
*LNCSOX17* cellular and molecular characterization. **(A)** Time resolved qRT-PCR profiling *SOX17* (green) and *LNCSOX17* (orange) transcript levels during endoderm differentiation (normalized to the housekeeping gene *18S*). Symbols indicate the mean and error bars indicate SD across three independent experiments. **(B)** Lineage tree heatmap showing *SOX17* (green) and *LNCNSOX17* (orange) expression across EN derived embryonic and adult tissues as measured by RNA-seq, derived from a curated data set of the Roadmap Epigenome Project(*140*) (Table 1). TPM, transcripts per million. aPS, anterior primitive streak; AFE, anterior foregut endoderm; PFE, posterior foregut endoderm; MHG; mid-hindgut; PPT, Peyer’s patch tissue; S, sigmoid; T, transverse. **(C)** smRNA-FISH of *LNCSOX17* in PSCs (left) and EN cells (right) counter-stained with Hoechst. Red arrowheads indicate two brighter and bigger foci present in each cell, potentially representing sites of nascent transcription. Scale bars, 10µm. **(D)** Frequencies of *LNCSOX17* smRNA-FISH foci in the nuclear (grey) or the cytoplasmic (white) compartments. *n*=79, number of analyzed cells. Lines of the violin plot indicate interquartile range around the median value. In the stacked barplot, error bars indicate SD around the mean value. **(E)** Barplots showing coding potential scores of randomly sampled *LNCRNA* ORFs (*n*=257,992) (grey) versus *LNCSOX17* ORFs (*n*=40) (orange). Scores are shown on the x-axis while ORF-density is plotted on the y-axis. Both conditions area is equal and compared to *SOX17* ORFs as coding gene control. n, number of analyzed ORFs. **(F)** Schematic of *LNCSOX17* isoform structure constructed from MinIONseq reads of endoderm cDNA. Exons are shown in orange while poly(A) is shown in white. The arrow indicates the transcriptional start site (TSS). Pie chart shows isoform reads (Ex1+2 black n=16, Ex1+3 grey n=11) and “sloppy spliced” (white *n*=89) transcript distribution as measured by MinIONseq (Table 1).

We further investigated *LNCSOX17* localization by single-molecule RNA fluorescence *in situ* hybridization (smRNA-FISH) and found it highly enriched at foci within the nuclear compartment, a characteristic feature of non-coding transcripts (median of 40 foci/cell, Figure 2C,D). Although we already named it as a non-coding RNA, we needed to assess the coding potential of *LNCSOX17* and used PhyloCSF to show that 37 of 40 predicted *LNCSOX17* open reading frames (ORFs) would likely result in no functional protein (Figure 2E). This is comparable to other short ORFs (sORFs) in the human lncRNA catalog (Figure 2E) (*45*). Notably, even the coding potential of the remaining three sORFs is about two orders of magnitude lower than for the *SOX17* coding sequence (Figure 2E).

To explore the structure and splicing variants of *LNCSOX17,* we used long-read Nanopore sequencing of definitive endoderm cDNA. The two most prevalent isoforms account for 23.3% of the split-reads, while 76.7% appear inconsistently spliced; a feature, which is frequently observed in lncRNAs (*46–50*) (termed ‘sloppy’ splicing, Figure 2F - Figure supplement 2F). Additionally, we used 5’ and 3’ rapid amplification of cDNA end (RACE) to determine the exact transcriptional start and end sites as well as the corresponding polyadenylation signal (Figure 2F - Figure supplement 2G). Taken together, our results show that the early definitive endoderm specific 22 kb lincRNA creates a ‘sloppily spliced’ transcript localized within the nuclear compartment.

### *LNCSOX17* does not regulate SOX17

To investigate the functional role of *LNCSOX17* during endoderm formation, we first generated a cell line carrying a constitutive transcriptional repressor (dCas9-KRAB-MeCP2 (*51*)) and then derived two clonal cell lines from it, harboring either a sgRNA targeting a control (sgCtrl), derived from a randomization approach of human TSS regions (*52*), or the *LNCSOX17* promoter (sgLNCSOX17) (see Methods) (Figure 3A). Immunofluorescent staining for dCas9-KRAB-MeCP2 demonstrated its homogeneous expression in the parental cell line (Figure supplement 3A). The dCas9 mediated silencing resulted in a strong repression of *LNCSOX17* RNA compared to the control, which was further validated by smRNA-FISH (Figure 3B,C - Figure supplement 3B). Functional validation of our repression system revealed H3K9me3 enrichment around the *LNCSOX17* promoter in sgLNCSOX17 cells, with a certain degree of spreading towards the enhancer *eSOX17* but no apparent consequence on *SOX17* regulation (Figure supplement 3C). To assess possible effects on *SOX17* further, we performed Capture Hi-C (cHi-C) in both cell lines. We could not observe any significant interaction differences (Log_2_FC = 0.02 p = 0.049) between the two cell lines within the *SOX17*-loop domain in definitive endoderm (Figure supplement 3D). Nevertheless, virtual 4C analysis revealed a marginal decrease in the *SOX17* enhancer-promoter interaction in the absence of *LNCSOX17* (Figure 3D). Despite this topological difference, loss of *LNCSOX17* does not appear to affect *SOX17* transcriptional activation and expression levels, indicating preserved enhancer functionality (Figure 3D,E).

**Figure 3:**
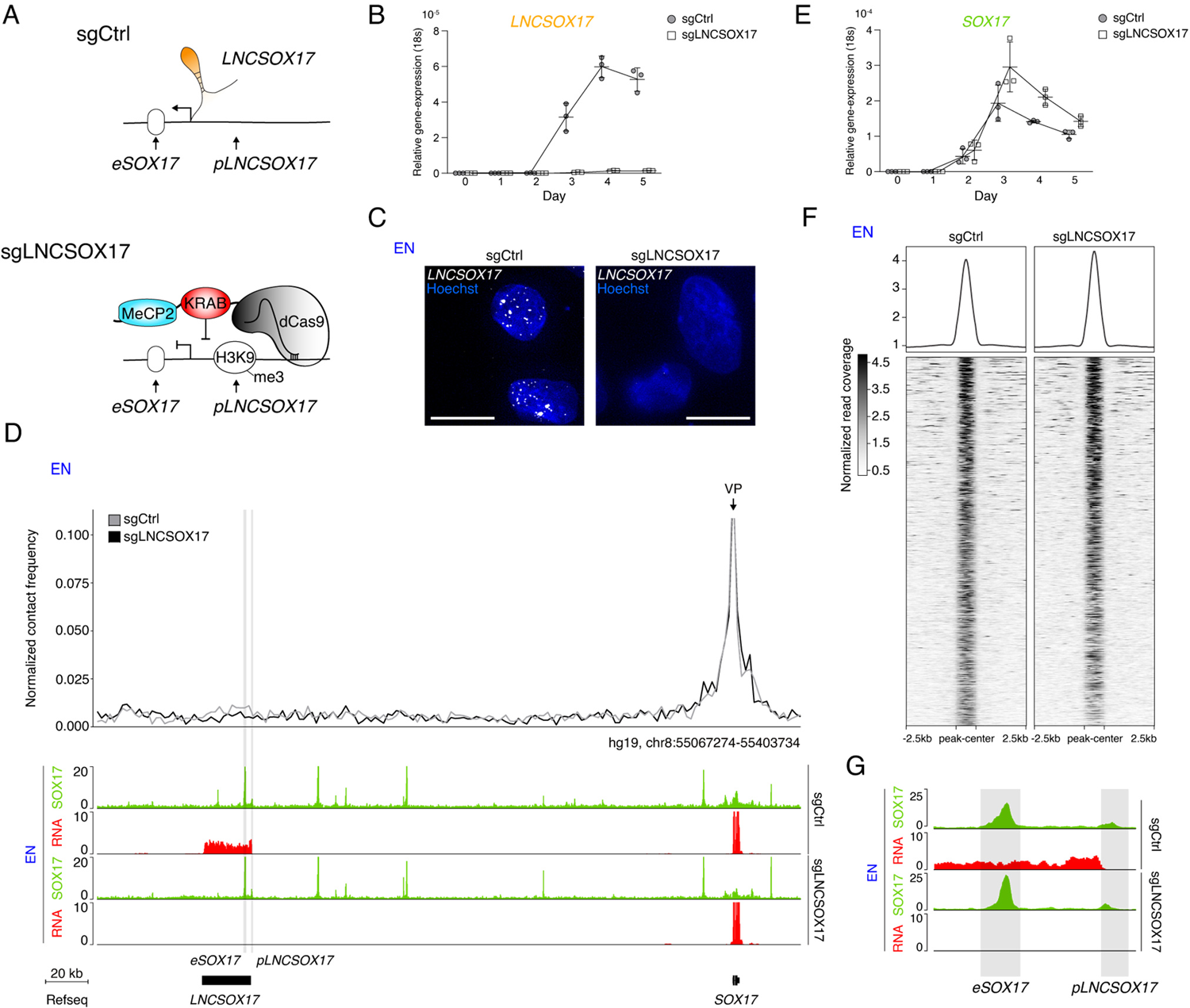
*LNCSOX17* regulation at the *SOX17* topological domain. **(A)** Schematic of *LNCSOX17* locus regulation in the absence (top) or presence (bottom) of a targeting dCas9-KRAB-MeCP2 complex, decorating *LNCSOX17* promoter with an H3K9me3 mark 355 bp upstream of the TSS. **(B)** Time-resolved qRT-PCR showing the expression of *LNCSOX17* during EN differentiation in the presence or absence of dCas9-KRAB-MeCP2 complex targeting *LNCSOX17* promoter (normalized to the housekeeping gene *18S*). Symbols indicate the mean and error bars indicate SD across three independent experiments. Individual data points are displayed. **(C)** smRNA-FISH of *LNCSOX17* in sgCtrl (left) and sgLNCSOX17 (right) EN cells counter-stained with Hoechst. Scale bars, 10 µm. For an extended field of view see Figure supplement 3B. **(D)** Virtual 4C analysis from capture Hi-C experiments in sgCtrl and sgLNCSOX17 EN cells using *SOX17* promoter as viewpoint, with 2kb resolution (upper panel). SOX17 EN ChIP-seq (RPKM) and RNA-seq (CPM) profiles in the two conditions are shown in the tracks (lower panel). *eSOX17* and *pLNCSOX17* are highlighted in grey. **(E)** Time-resolved qRT-PCR showing the expression of *SOX17* during EN differentiation in the presence or absence of dCas9-KRAB-MeCP2 complex targeting *LNCSOX17* promoter (normalized to the housekeeping gene *18S*). Symbols indicate the mean and error bars indicate SD across three independent experiments. Individual data points are displayed. **(F)** Heatmap showing SOX17 binding distribution genome-wide in sgCtrl and sgLNCSOX17 EN. The displayed peaks represent the union of the identified peaks in the two conditions (n=61694). **(G)** SOX17 ChIP-seq and RNA-seq tracks at the *LNCSOX17* locus showing SOX17 binding at the SOX17 enhancer (*eSOX17*) and LNCSOX17 promoter (*pLNCSOX17*). SOX17 binding on *pLNCSOX17* results in *LNCSOX17* activation, if *pLNCSOX17* is not targeted by dCas9-KRAB-MeCP2.

Next, we performed SOX17 Chromatin Immunoprecipitation sequencing (ChIP-seq) in our control and loss of function cell line. This demonstrated that there is no change in SOX17 occupancy at the *SOX17* locus as well as genome-wide occupancy despite the loss of *LNCSOX17* (Figure 3D,F). Interestingly, we found SOX17 enrichment at the *LNCSOX17* promoter (*pLNCSOX17*), potentially contributing to its activation – consistent with the timed *LNCSOX17* activation relative to SOX17 (Figure 3D,G and 2A). To further explore the regulation of *LNCSOX17* by SOX17, we performed sgRNA/Cas9 mediated *SOX17* gene ablation, retrieving heterozygous (*SOX17^WT/Δ^*) and homozygous (*SOX17^Δ/Δ^*) knock out cell lines (Figure supplement 3E-G). Notably, SOX17 disrupted cells fail to induce *LNCSOX17* (Figure supplement 3H).

In order to distinguish between the function of *LNCSOX17* active transcription and its actual transcript (*29, 53*), we generated another cell line by introducing a strong transcriptional termination signal downstream of an mRuby cassette in the first exon of *LNCSOX17,* hereafter LNCSOX17^p(A)/p(A)^ (Figure supplement 3I-K). qRT-PCR demonstrated that the expression of *LNCSOX17* is abolished in LNCSOX17^p(A)/p(A)^ EN cells, while the mRuby cassette is actively transcribed, indicating ongoing transcription at the locus in an endoderm specific manner (Figure supplement 3L). In line with our depletion experiments, SOX17 levels are not affected in LNCSOX17^p(A)/p(A)^ EN cells (Figure supplement 3L). These results clearly demonstrate that *LNCSOX17* induction is dependent on SOX17, whereas the *LNCSOX17* transcript and the act of transcription are dispensable for SOX17 activation as well as its genome-wide localization.

### *LNCSOX17* folds into secondary structures and interacts with HNRNPU

To explore what and how *LNCSOX17* is involved in endoderm regulation, we investigated whether it had the sequence characteristic necessary to fold into secondary structures containing stretches of double-stranded RNA, which are often recognized by RNA binding proteins and a common way lncRNAs exert their functions (*54–58*). Both *LNCSOX17* isoforms are predicted to fold into secondary structures containing stem-loops and double-stranded RNA regions with high confidence (Figure supplement 4A). To identify possible protein interaction partners, we performed RNA-pulldown followed by mass spectrometry (Figure supplement 4B-C). Among the putative *LNCSOX17* interactors, we identified several heterogenous nuclear ribonucleoprotein (hnRNP) family members, including HNRNPU (Figure supplement 4D). HNRNPU is a ribonucleoprotein previously reported to interact with lncRNAs to regulate various functions during development including nuclear matrix organization (*17, 59*), X chromosome inactivation (*60*), RNA splicing (*61, 62*), and epigenetic control of gene expression (*63–65*). To validate HNRNPU-*LNCSOX17* interaction, we performed HNRNPU RNA immunoprecipitation (RIP) (Figure supplement 4E,F). We found *LNCSOX17* to be enriched in the HNRNPU immunoprecipitation, to levels comparable to known RNA interactors such as *XIST* or *NEAT1* (Figure supplement 4G).

Although more work is required, our preliminary analysis identified the known lncRNA-interacting ribonucleoprotein HNRNPU as a *LNCSOX17* partner and indicates a possible route for its molecular function.

### *LNCSOX17* is required for the differentiation towards definitive endoderm

To investigate the cellular role of *LNCSOX17*, we performed immunofluorescent staining and fluorescent activated cell sorting (FACS) for CXCR4 in control and *LNCSOX17* depleted cells. The latter showed a substantial reduction in the CXCR4^+^ cell population during differentiation, suggesting hampered differentiation potential towards endoderm (Figure 4A). However, consistent with the transcriptional data, SOX17 protein levels were not affected (Figure 4A). Both phenotypes were recapitulated in the LNCSOX17^p(A)/p(A)^ EN cells (Figure supplement 5A). As expected, based on its highly restricted expression, differentiation towards the other two germ layers (mesoderm and ectoderm) was not affected (Figure supplement 5B,C and 2D).

**Figure 4:**
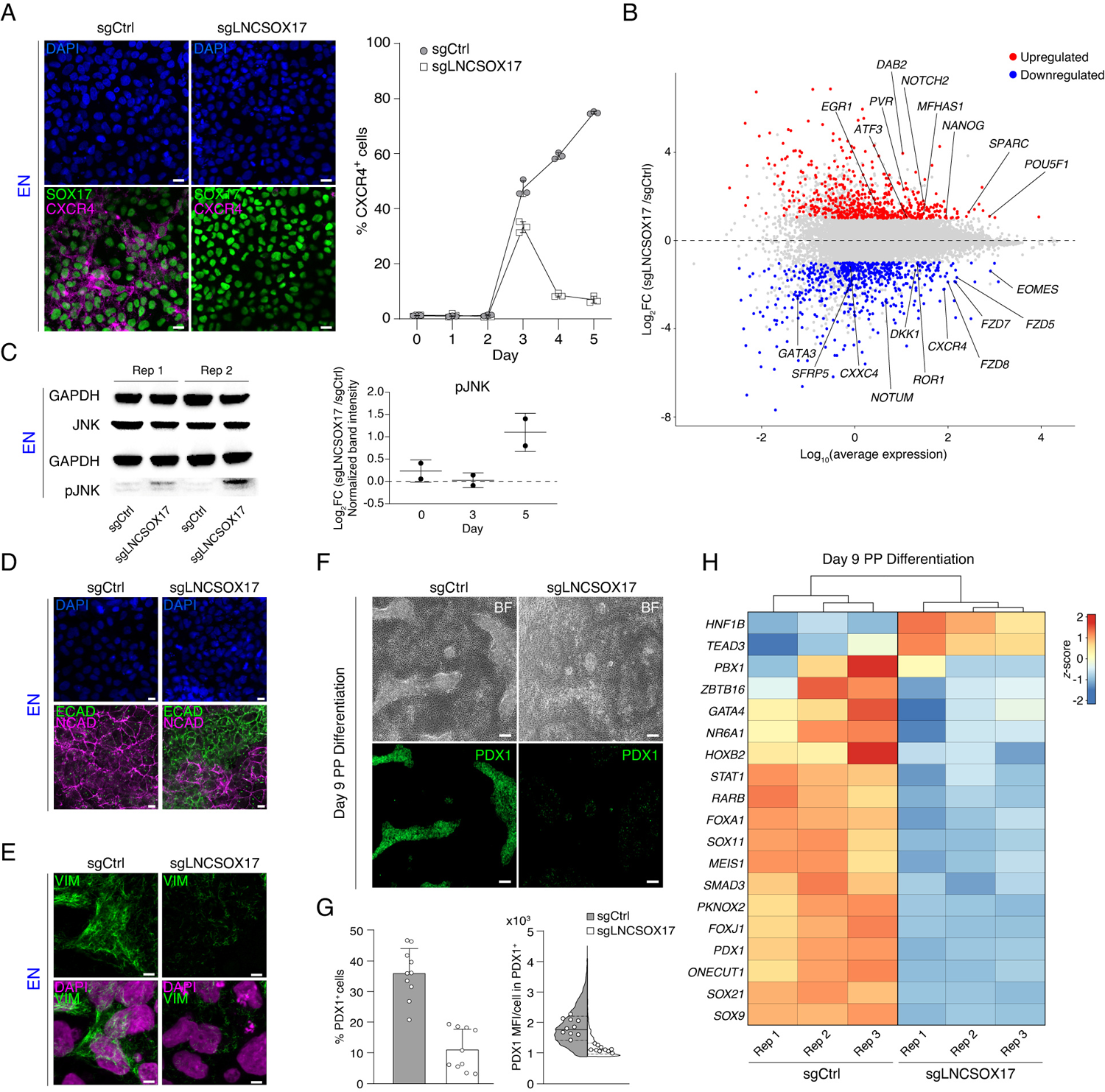
Endodermal defects in cells depleted of *LNCSOX17*. **(A)** Immunofluorescent (IF) staining of SOX17 and CXCR4 in EN cells expressing either sgCtrl or sgLNCSOX17 counter-stained with DAPI (left panel). Line plot showing percentage of FACS-derived CXCR4^+^ cell-population at given time-points during endoderm differentiation (right panel). Symbols indicate mean values, while error bars show SD across three independent experiments. Individual data points are displayed. Scale bars, 10 µm. **(B)** Scatter plot highlighting differentially expressed genes between sgLNCSOX17 and sgCtrl EN cells. Significantly (Log_2_FC ≥ 1) upregulated genes (*n*=590) upon *LNCSOX17* repression are shown in red while significantly (Log_2_FC ≤ −1) down-regulated genes (n=584) are shown in blue. The complete lists of TPMs and differentially expressed genes are provided in Table 2. **(C)** JNK and pJNK Western Blots of sgCtrl and sgLNCSOX17 EN cells (left panel). GAPDH signals are used as loading controls above the corresponding JNK/pJNK signals. Boxplot showing relative pJNK levels during endoderm differentiation shown as Log_2_FC of sgLNCSOX17 over sgCtrl (right panel). Central line indicates the mean, error bars indicate the SD across two independent experiments. Differentiation time-course blot is shown in Figure supplement 5G. **(D)** IF staining of ECAD and NCAD in EN cells expressing either sgCtrl or sgLNCSOX17 counter-stained with DAPI. Scale bars, 10µm. **(E)** IF staining of VIM in EN cells expressing either sgCtrl or sgLNCSOX17 counter-stained with DAPI. Scale bars, 5µm. **(F)** Bright field images of PP differentiation cultures (upper panel) followed by IF staining for PDX1 (lower panel) of either sgCtrl or sgLNCSOX17 cells. Scale bars, 10 µm. **(G)** IF staining quantification of overall (sgCtrl, *n*=17657, sgLNCSOX17, *n*=5279) PDX1^+^ population percentages (left) or PDX1 mean fluorescence intensity distribution in PDX1^+^ cells (right). Bar plot error bars indicate SD around the mean value and white dots represent mean values for the individual replicates (N=10). Lines of the violin plot indicate interquartile range around the median value and white dots represent median values for the individual replicates (N=10). List of value for each cell and corresponding statistics are shown in (Table 4). **(H)** Heatmap showing row-normalized *z*-scores of PP specific marker genes(*80*) in sgCtrl and sgLNCSOX17 EN cells as measured by RNA-seq at day 9 of differentiation. Columns were ordered by hierarchical clustering (represented as tree above the heatmap). Note the reduced expression of PP master transcription factor *PDX1* in sgLNCSOX17 as compared to sgCtrl. The complete lists of TPMs and differentially expressed genes are provided in Table 2.

To characterize the differentiation defect on a molecular level, we performed time-resolved RNA-seq in *LNCSOX17* depleted and control cell lines on day 0, 3, and 5 of endoderm differentiation. Principal Component Analysis (PCA) revealed only marginal variance by day 3, while a more substantial transcriptional divergence was observed on day 5 (Figure supplement 5D). Differential gene expression analysis identified 584 significantly down- and 590 significantly upregulated genes in *LNCSOX17* depleted cells at day 5 (Figure 4B). In particular, we found pluripotency genes (e.g., *POU5F1*, *NANOG*) and endoderm/WNT related genes (e.g., *EOMES*, *GATA3*, *CXCR4, FZD5*, *FZD7*, *FZD8*, *DKK1*, *NOTUM*, *ROR1*, *CXXC4*, *SFRP5)* to be significantly up- and downregulated, respectively (Figure 4B - Figure supplement 5E). Time resolved qPCR analysis over 5 days confirmed, a lack of key endoderm marker activation and expression in *LNCSOX17* depleted cells (including *CXCR4, GATA3, GATA4, KLF5, CPE, GPR, HHEX, EPSTI1, FOXA3*), an aberrant transcriptional signature we observe also in LNCSOX17^p(A)/p(A)^ EN cells (*35, 66–71*) (Figure supplement 5F-H). Interestingly, among the significantly, upregulated genes in *LNCSOX17* depleted cells, we found an enrichment of *JUN* (AP-1) pathway target genes (including *EGR1*, *ATF3*, *PVR*, *DAB2*, *NOTCH2*, *MFHAS1*, *SPARC*) (*72–77*), which has recently been described to act as a barrier for the exit from pluripotency towards endoderm formation (Figure 4B - Figure supplement 5E) (*32*). Phosphorylation levels of JUN-activating upstream kinase JNK are a strong indicator of JUN pathway activation (*32, 78, 79*), which we observed by increased relative amounts of pJNK in *LNCSOX17* depleted cells (Figure 4C - Figure supplement 6A). Inhibition of JNK hyperactivity (JNK Inhibitor XVI) from day 3 of definitive endoderm differentiation only partially rescued the specification defect in *LNCSOX17* depleted cells (Figure supplement 6B,C).

Furthermore, immunofluorescent staining for ECAD, NCAD, and VIM revealed retention of an epithelial signature in *LNCSOX17* depleted endoderm cells (Figure 4D,E - Figure supplement 5E and 6D,E). Moreover, VIM-signal distribution within *LNCSOX17* depleted cells was also altered, indicating a potential cellular polarization defect (Figure 4E - Figure supplement 6E).

Finally, we evaluated if these cells have lost the potential to further differentiate into pancreatic progenitor (PP) cells (*80*). Immunofluorescent staining identified a very distinct PDX1^+^ population in the control cells after nine days of directed differentiation, which is notably reduced in *LNCSOX17* depleted cells (Figure 4F,G - Figure supplement 6F). In addition, transcriptomic analysis of differentiated control and *LNCSOX17* depleted cells indicates a substantial gene expression difference, including the specific downregulation of pancreatic progenitor marker genes (*80*) (Figure 4H - Figure supplement 6G). Our data therefore highlight the importance of *LNCSOX17* for the induction of definitive endoderm and subsequent differentiation potential.

## DISCUSSION

Here, we describe the discovery and characterization of *LNCSOX17* as a functionally essential lincRNA in human definitive endoderm. Most lncRNAs act locally, regulating the chromatin architecture and the expression of neighboring genes in *cis* (*81–84*), especially when overlapping with enhancer elements. In particular, the fine-tuned expression of several developmental transcription factors has been shown to rely on the activity of lncRNAs present within the same topological domain. (*25, 82, 85*). Interestingly, *LNCSOX17* appears distinct from these and other endodermal specific lncRNAs (*27, 29, 86*) as it does not appear to regulate the adjacent *SOX17* gene. The use of two orthogonal loss of function approaches in our work (suppression of *LNCSOX17* activation and early termination) showed that *LNCSOX17* transcription is dispensable for proper *SOX17* regulation. It remains to be determined what the targets and regulatory mechanism of *LNCSOX17* are. One may speculate that these could also be distant and unrelated loci to the *SOX17* loop-domain, as we find many *LNCSOX17* distinct puncta in the nuclear compartment of endodermal cells. Typically, local *cis*-acting lncRNAs mainly show accumulation at the two sites of nascent transcription (*27, 29, 30, 87*). The observed interaction with the HNRNP complex may link it to various nuclear-related functions needed for endoderm specification. It remains to be determined how this compares to other endodermal lncRNAs, which mainly exert their functions together with endoderm-specific transcription factors (*27, 28, 30, 88*).

SOX genes are fundamental transcription factors that have a variety of functions including the specification of cell types and tissues during embryonic development. They are evolutionary conserved and evolved as a result of a series of ancient genomic duplication events (*89*). Interestingly, at other SOX gene loci, the presence of one or multiple lncRNAs have been reported, but these lncRNAs, in contrast to *LNCSOX17*, are involved in the modulation of the associated SOX gene expression in *cis* (*90–93*). This suggests that lncRNAs near paralogous genes may evolve distinct role and regulatory mechanisms.

At a functional level, our results show that *LNCSOX17* is essential for definitive endoderm specification and its loss limits further downstream differentiation, as demonstrated by the pancreatic progenitor differentiation. How the different phenotypic changes associated with the loss of *LNCSOX17* arise, such as an aberrant endodermal transcriptome, EMT-failure, JNK-hyperactivity and lack of pancreatic progeny, remains unclear. Similar to other lncRNAs, the function of *LNCSOX17* could depend on specific protein interactors during endodermal fate specification. Biochemical assays to simultaneously profile RNA-RNA, RNA-DNA and RNA-protein interactions (*94–96*) might help elucidating the precise mechanism of action by which *LNCSOX17* controls endodermal transition. From the developmental perspective, *LNCSOX17* and its transient, highly stage-specific nature make it an intriguing regulator compared to most of the protein-coding genes, including endodermal transcription factors, e.g. SOX17, FOXA2 and GATA4, which are expressed longer and in a variety of somatic tissues. In this context, it is worth noting that the development of definitive endoderm during human gastrulation *in vivo* takes place within hours and the gene regulatory network (GRN) governing this transition has to be tightly controlled (*33, 39, 97*). Given our strong phenotype and the detection of its expression in the gastrulating human embryo, it is possible that this lncRNA and others play crucial roles during such developmental windows.

As such, our study provides an important piece towards a more complete understanding of the multi-layered complex regulation of human cellular differentiation, especially human definitive endoderm specification, and connects it to a previously unannotated non-coding RNA. Overall, in our study we identified and characterized a previously unknown lncRNA that we termed *LNCSOX17* and demonstrate its critical role in the formation of human definitive endoderm. *LNCSOX17* is highly tissue-specific, spliced and mainly localized in the nucleus. Despite its sequence overlapping with SOX17 regulatory element, *LNCSOX17* does not regulate SOX17 within the topological domain. As such our study finds a new non-coding RNA that broadens the range of non-coding mechanisms and our general understanding of RNA based nuclear regulation.

## MATERIAL AND METHODS

Default parameters were used, if not otherwise specified, for all software and pipelines utilized in this study.

### 1. Molecular cloning of *SOX17* and e*SOX17.2* knock-out constructs

For CRISPR/Cas9 mediated targeting of either *SOX17* or *eSOX17.2* we utilized pSpCas9(BB)-2A-GFP (*98*) (PX458), which was a gift from Feng Zhang (Addgene plasmid # 48138; http://n2t.net/addgene:48138; RRID:Addgene_48138) (*98*). Prior to small guide RNA (sgRNA) cloning, PX458 was initially modified and further re-named into P2X458. P2X458 harbors an additional independent U6-promoter followed by a small guide RNA (sgRNA) scaffold expression cassette, which allows the insertion of an additional sgRNA by SapI restriction enzyme cloning. To generate P2X458, PX458 and the synthetized SapI sgRNA expression cassette (IDT, find sequence in Table 3) were digested with KpnI (New England Biolabs, R3142S). Next, the SapI sgRNA expression cassette was ligated into the KpnI linearized PX458 in a 3:1 molarity ratio using T4 DNA-ligase (New England Biolabs, M0202S) according to the manufacturer’s instructions followed by transformation and Sanger sequencing to verify successful cloning. sgRNA-cloning was performed with NEBuilder HiFi DNA Assembly Master Mix (New England Biolabs, E2621S) according to manufacturer’s instructions using BbsI-linearization of P2X458 for the first sgRNA and SapI linearization of P2X458 for the second sgRNA as backbone, combined with single stranded oligonucleotides containing the sgRNA sequences as inserts (1:3 molar ratio) (find sequence in Table 3). Bacterial transformation and Sanger sequencing were performed to verify successful cloning.

### 2. Molecular cloning of Luciferase reporter constructs

pGL4.27[luc2P/minP/Hygro] (Promega, E8451) containing a minimal CMV-promoter for enhancer-assays or pGL4.15[luc2P/Hygro] (Promega, E6701) w/o any promoter for promoter-assays were first digested using EcoRV (New England Biolabs, R3195S). Next, full *eSOX17*, *eSOX17.1* or *eSOX17.2* for enhancer-assays and *pSOX17* or *pLNCSOX17* genomic regions were PCR amplified with primers containing homology overhangs to the plasmid. PCR products were purified and cloned into the linearized plasmid utilizing the NEBuilder HiFi DNA Assembly Master Mix (1:3 molar ratio) according to the manufacturer’s instructions. Bacterial transformation followed by Sanger sequencing verified the successful cloning. Cloning primers are listed in (Table 3).

### 3. Molecular cloning of lentiviral sgRNA constructs

pU6-sgRNA EF1Alpha-puro-T2A-BFP (*52*) was digested with BstXI (New England Biolabs, R0113S) and BlpI (New England Biolabs, R0585S) and the linearized plasmid was gel extracted with the QIAquick Gel Extraction Kit (Quiagen, 28704). Subsequently sgRNAs (sgLNCSOX17 or sgCtrl) (s. Table 3) were cloned in the linearized backbone using NEBuilder HiFi DNA Assembly Master Mix (1:3 molar ratio) according to the manufacturer’s instructions. Bacterial transformation and sanger sequencing confirmed the successful cloning. pU6-sgRNA EF1Alpha-puro-T2A-BFP was a gift from Jonathan Weissman (Addgene plasmid # 60955; http://n2t.net/addgene:60955; RRID:Addgene_60955).

### 4. Molecular cloning of *SOX17* reporter knock-in constructs

pUC19 plasmid was digested with SmaI (New England Biolabs, R0141S) and the linearized plasmid was gel extracted with the QIAquick Gel Extraction Kit (Quiagen, 28704). Next, *SOX17* homology arm genomic regions were PCR amplified with primers containing homology overhangs to the plasmid and to a T2A-H2B-mCitrine-loxP-hPGK-BSD-loxP selection cassette.

The left homology arm overlapped with the end of the *SOX17* coding sequence, and the T2A-H2B-mCitrine cassette which was cloned in frame with the last *SOX17* aminoacid. PCR products and selection cassette were purified and cloned into the linearized plasmid utilizing the NEBuilder HiFi DNA Assembly Master Mix according to the manufacturer’s instructions. Bacterial transformation followed by Sanger sequencing verified the successful cloning.

sgRNA targeting the genomic region of integration was cloned in BbsI linearized pX335-U6-Chimeric_BB-CBh-hSpCas9n (*99*) (D10A) plasmid (Addgene #42335) using NEBuilder HiFi DNA Assembly Master Mix (1:3 molar ratio) according to the manufacturer’s instructions. pX335-U6-Chimeric_BB-CBh-hSpCas9n(D10A) was a gift from Feng Zhang (Addgene plasmid # 42335; http://n2t.net/addgene:42335; RRID:Addgene_42335) Bacterial transformation and sanger sequencing confirmed the successful cloning. Cloning primers are listed in (Table 3).

### 5. Molecular cloning of *LNCSOX17*-promoter-KI constructs

pUC19 plasmid was digested with SmaI (New England Biolabs, R0141S) and the linearized plasmid was gel extracted with the QIAquick Gel Extraction Kit (Quiagen, 28704). Next, *LNCSOX17* homology arm genomic regions were PCR amplified with primers containing homology overhangs to the plasmid and to a mRuby-3xFLAG-NLS-3xSV40-poly(A)-loxP-mPGK-PuroR-loxP selection cassette.

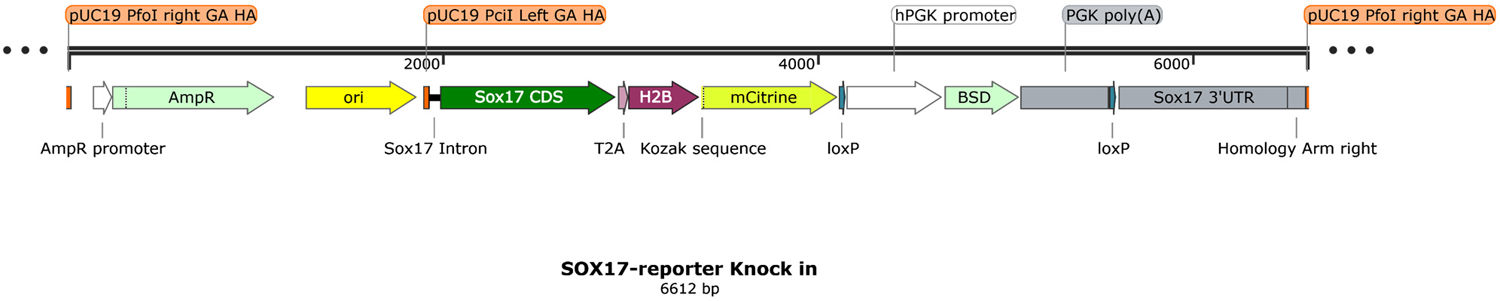

The left homology arm overlapped with the *LNCSOX17* promoter including 30 bp of *LNCSOX17* Exon 1, and a mRuby-3xFLAG-NLS-3xSV40-poly(A) cassette which was cloned +30 bp after *LNCSOX17*-TSS into Exon 1. The right homology arm overlapped with *LNCSOX17* Exon 1-30 bp TSS, and a loxP-mPGK-PuroR-loxP cassette which was cloned following the mRuby-3xFLAG-NLS-3xSV40-poly(A) cassette. Both, the mRuby-3xFLAG-NLS-3xSV40-poly(A) and the loxP-mPGK-PuroR-loxP cassette also shared homology. All PCR products were purified and cloned into the linearized plasmid utilizing the NEBuilder HiFi DNA Assembly Master Mix according to the manufacturer’s instructions. Bacterial transformation followed by Sanger sequencing verified the successful cloning.

For CRISPR/Cas9 mediated targeting of the *LNCSOX17* promoter we utilized pSpCas9(BB)-2A-GFP (*98*) (PX458), which was a gift from Feng Zhang (Addgene plasmid # 62988; http://n2t.net/addgene:62988; RRID:Addgene_62988) (*98*). sgRNA-cloning was performed with NEBuilder HiFi DNA Assembly Master Mix (New England Biolabs, E2621S) according to manufacturer’s instructions using BbsI-linearization of PX458, combined with single stranded oligonucleotides containing the sgRNA sequences as inserts (1:3 molar ratio) (find sequence in Table 3). Bacterial transformation and Sanger sequencing were performed to verify successful cloning.

### 6. hiPS cell culture

ZIP13K2 (*100*) hiPSCs were maintained in mTeSR1 (Stemcell Technologies, 85850) on pre-coated culture ware (1:100 diluted Matrigel (Corning, 354234) in KnockOut DMEM (Thermo Fisher Scientific, 10829-018)). Clump-based cell splitting was performed by incubating the cells in final 5 mM EDTA pH 8,0 (Thermo Fisher Scientific, 15575-038) in DPBS (Thermo Fisher Scientific, 14190250) 5 min at 37°C, 5% CO_2_. Single cell splitting was performed by incubating the cells with Accutase (Sigma-Aldrich, A6964) supplemented with 10 µM Y-27632 (Tocris, 1254) for 15 min at 37°C, 5% CO_2_. Cell counting was performed using a 1:1 diluted single-cell suspensions in 0,4% Trypan Blue staining-solution (Thermo Fisher Scientific, 15250061) on the Countess II automated cell-counter (Thermo Fisher Scientific). Wash-steps were performed by spinning cell-suspensions at 300 x g 5 min at room temperature (RT).

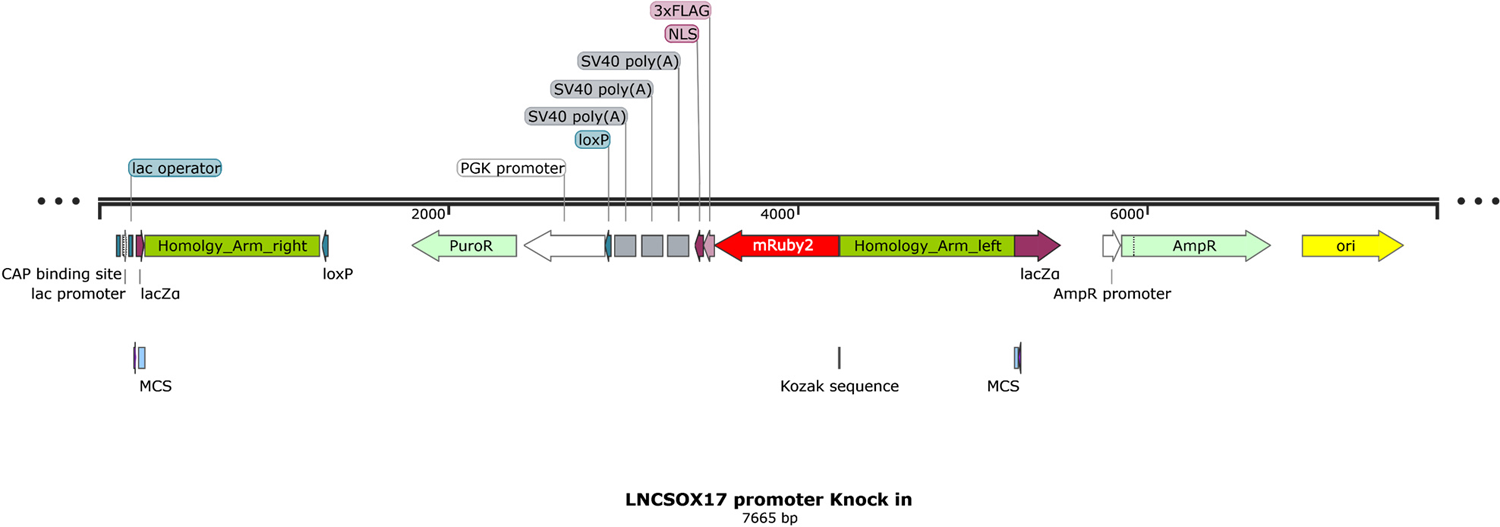

### 7. Definitive endoderm (EN) differentiation

To guarantee high reproducibility, constant media-quality, and mTeSR1 compatibility, definitive endoderm differentiations were exclusively performed utilizing the STEMdiff Trilineage Endoderm Differentiation media (Stemcell Technologies, 05230). Single cell suspensions of mTeSR1 maintained ZIP13K2 hiPSCs were seeded into the respective culture formats according to the required cell-number as recommended by the manufacturer’s instructions. Media change using the STEMdiff Trilineage Endoderm Differentiation media was performed on a daily bases according to the manufacturer’s instructions. Cells were then collected at required timepoints by washing the plate with DPBS before single-cell dissociation was performed with Accutase for 15 min at 37°C, 5% CO_2_. Single cell suspensions of definitive endoderm (EN) differentiated cells were utilized for further downstream analysis (qPCR, Western Blot, FACS etc.).

### 8. Embryoid body (EB) formation followed by ScoreCard Assay

ZIP13K2 hiPSC single cell suspensions were prepared and counted as previously described (s. *hiPS cell culture*). Next, 1 x 10^3^ cells/well of either sgCtrl or sgLNCSOX17 hiPSCs were seeded on a 96-well ultra-low attachment U-bottom plate (Corning, 7007) in respective cell culture media.

#### Random EB differentiation

Cells were seeded in 200 µl /well of hES-media (Final DMEM-F12 (Thermo Fisher Scientific, 11320074), 20% KSR (Thermo Fisher Scientific, 10828028), 1% Penicillin /Streptomycin, 1% NEAA (Thermo Fisher Scientific, 11140050), 0,5% GlutaMAX, HEPES (Thermo Fisher Scientific, 31330038)), supplemented with final 10 µM Y-27632. Single cell suspensions were spun at 100 x g for 1 min at RT and further cultured for 16 h at 37°C, 5% CO_2_. The following day 150 µl media supernatant was carefully exchanged by 150 µl fresh hES-media (without Y-27632). Cells were further cultured for additional 48 h at 37°C, 5% CO_2_. The very same media was replaced every 48h until day 9. At day 9, EBs were collected, washed once in DPBS and RNA isolated (s. *RNA isolation and cDNA synthesis*).

#### Undifferentiated control EBs

Cells were seeded in 200 µl /well of mTeSR1, supplemented with final 10 µM Y-27632. Single cell suspensions were spun at 100 x g for 1 min at RT and further cultured for 16 h at 37°C, 5% CO_2_. The following day 150 µl media supernatant was carefully exchanged by 150 µl fresh mTeSR1 media (without Y-27632). Cells were further cultured for additional 48 h 37°C, 5% CO_2_. At day 3, EBs were collected, washed once in DPBS and RNA isolated (s. *RNA isolation and cDNA synthesis*). cDNA-conversion and ScoreCard assay (Thermo Fisher Scientific, A15870) has been performed according to the manufacturer’s instructions.

### 9. JNK inhibition experiments

For the JNK-inhibition experiments, 1 µM JNK inhibitor XVI (Sellekchem, S4901) final was supplemented to the media from day 3 of EN differentiation onward. The corresponding volume of DMSO was supplemented to the media of the control samples.

### 10. Pancreatic progenitor (PP) differentiation

Pancreatic progenitor (PP) differentiation was performed as previously described (*80*) with minor changes. Briefly, single cell suspensions of ZIP13K2 hiPSCs (s. *hiPS cell culture*) were seeded at a density of 5 x 10^5^ cells /cm^2^ in mTeSR1 supplemented with 10 µM Y-27632. After 24 h, culture medium was replaced with S1-media (Final 11,6 g/L MCDB131, Sigma Aldrich, M8537-1L; 2 mM D-+-Glucose, Sigma Aldrich, G7528-250G; 2,46 g/L NaHCO3, Sigma Aldrich, S5761-500G; 2% FAF-BSA, Proliant Biologicals, 68700-1; 1:50000 of 100 x ITS-X, Thermo Fisher Scientific, 51500056; 1 x GlutaMAX, Thermo Fisher Scientific, 35050-038; 0,25 mM ViatminC, Sigma-Aldrich, A4544-100G; 1% Pen-Strep, Thermo Fisher Scientific, 15140122) supplemented with final 100 ng/ml Activin-A (R&D Systems, 338-AC-01M) and 1,4 µg/ml CHIR99021 (Stemgent, 04-0004-10). The following 2 days, cells were cultured in S1-media supplemented with final 100 ng/ml Activin-A. Next, cells were cultured in S2-media (Final 11,6 g/L MCDB131; 2 mM D-+-Glucose; 1,23 g/L NaHCO3; 2% FAF-BSA; 1:50000 of 100 x ITS-X; 1 x GlutaMAX; 0,25 mM ViatminC; 1% Pen-Strep) supplemented with final 50 ng/ml KGF (Peprotech, 100-19-1MG) for 48 h. After these 48 h, cells were cultured in S3-media (Final 11,6 g/L MCDB131; 2 mM D-+-Glucose; 1,23 g/L NaHCO3; 2% FAF-BSA; 1:200 of 100 x ITS-X; 1 x GlutaMAX; 0,25 mM ViatminC; 1% Pen-Strep) supplemented with final 50 ng/ml KGF (Peprotech, 100-19-1MG), 200 nM LDN193189 (Sigma Aldrich, SML0559-5MG), 0,25 µM Sant-1 (Sigma Aldrich, S4572-5MG), 2 µM Retinoic Acid (Sigma Aldrich, R2625-50MG), 500 nM PDBU (Merck Millipore, 524390-5MG) and 10 µM Y-27632 for 24 h. Finally, cells were cultured in the previous S3-media composition w/o supplementation of LDN193189 for 24 h. Between daily media changes, cells were washed once with 1 x DPBS. Throughout the entire differentiation process, cells were cultured at 37°C, 5% CO_2_ in 100 µl media /cm^2^.

### 11. Luciferase reporter assays

ZIP13K2 hiPSCs (s. *hiPS cell culture*) were treated with Accutase containing 10 µM Y-27632 for 15 min, 37°C, 5% CO_2_ to obtain a single cell suspension. Cell suspensions were counted and seeded at a density of 10^5^ cells /cm^2^ in mTeSR1 supplemented with final 10 µM Y-27632. Sixteen hours later, cells were co-transfected with 15 fmol pRL-TK (Promega, E2241) and 150 fmol of either pGL4.27[luc2P/minP/Hygro] empty vector or pGL4.27[luc2P/minP/Hygro] containing either *eSOX17*, *eSOX17.1* or *eSOX17.2* utilizing Lipofectamin Stem Transfection Reagent (Thermo Fisher Scientific, STEM00003) following the manufacturer’s instructions. Transfection was performed in mTeSR1 containing 10 µM Y-27632 for 16 h at 37°C, 5% CO_2_. Subsequently, endoderm differentiation was initiated (day 0) using the STEMdiff Trilineage Endoderm Differentiation media. At day 0, 2, 3 or 5 of endoderm differentiation, cells were lysed and Renilla as well as Firefly Luciferase activity was measured using the Dual-Glo Luciferase Assay System (Promega, E2920) according to the manufacturer’s instructions. Raw values (Table 1) were measured on the GloMax-Multi Detection System (Promega).

### 12. Generation of *SOX17* and *eSOX17.2* CRISPR/Cas9 knock-out hiPSC lines

ZIP13K2 hiPSCs (s. *hiPS cell culture*) were treated with Accutase containing final 10 µM Y-27632 for 15 min at 37°C, 5% CO_2_ to obtain a single cell suspension. Cell suspensions were counted and seeded at a density of 1-2 x 10^5^ cells /cm^2^ in mTeSR1 supplemented with final 10 µM Y-27632. Cells were pre-cultured for 16 h at 37°C, 5% CO_2_ prior to transfection.

Cells were then transfected with 6 µg of P2X458 using Lipofectamin Stem Transfection Reagent according to the manufacturer’s instructions. GFP^+^ cells were FACS-sorted 16-24 h post-transection with the FACSAria II or the FACSAria Fusion (Beckton Dickinson) and seeded at a density of 0,5-1 x 10^3^ cells /cm^2^ in mTeSR1 supplemented with 10 µM Y-27632 to derive isogenic clones. Single-cell derived colonies were manually picked, and split half for maintenance in a well of a 96-well plate and half used for genotyping using the Phire Animal Tissue Direct PCR Kit (Thermo Fisher Scientific, F140WH) following manufacturer’s instructions. Genotyping primer are listed in (Table 3). Edited alleles were verified by cloning PCR-products into the pJET1.2 backbone (Thermo Fisher Scientific, K1232) according to the manufacturer’s instructions, followed by bacterial transformation and sanger sequencing.

### 13. Generation of *SOX17*-reporter hiPS cell line

ZIP13K2 hiPSCs (s. *hiPS cell culture*) were treated with Accutase containing final 10 µM Y-27632 for 15 min at 37°C, 5% CO_2_ to obtain a single cell suspension. Cell suspensions were counted and seeded at a density of 1-2 x 10^5^ cells /cm^2^ in mTeSR1 supplemented with final 10 µM Y-27632. Cells were pre-cultured for 16 h at 37°C, 5% CO_2_ prior to transfection.

The following day, cells were transfected using Lipofectamin Stem Transfection Reagent in fresh mTeSR1 supplemented with final 10 µM Y-27632 for 24 h at 37°C, 5% CO_2_. Transfection mixtures contained 3 µg of T2A-H2B-mCitrine-loxP-hPGK-BSD-loxP donor plasmid and 3 µg of PX335-SOX17 (1:1 molar ratio).

Two days post transfection, cells were selected with final 2 µg/ml Blasticidin-S-HCl (Thermo Fisher Scientific, A1113903) for 14 days at 37°C, 5% CO_2_. For the derivation of isogenic reporter cell lines, single-cell derived colonies were manually picked and expanded. Differentiation into EN followed by FACS analysis was used to confirm clones that were activating the reporter.

### 14. Generation of *LNCSOX17*-promoter-KI hiPS cell line

ZIP13K2 SOX17-reporter (s. *Generation of SOX17-reporter hiPS cell line*) hiPSCs (s. *hiPS cell culture*) were treated with Accutase containing final 10 µM Y-27632 for 15 min at 37°C, 5% CO_2_ to obtain a single cell suspension. Cell suspensions were counted and seeded at a density of 1-2 x 10^5^ cells /cm^2^ in mTeSR1 supplemented with final 10 µM Y-27632. Cells were pre-cultured for 16 h at 37°C, 5% CO_2_ prior to transfection.

The following day, cells were transfected using Lipofectamin Stem Transfection Reagent in fresh mTeSR1 supplemented with final 10 µM Y-27632 for 24 h at 37°C, 5% CO_2_. Transfection mixtures contained 3 µg of mRuby-3xFLAG-NLS-3xSV40-poly(A)-loxP-mPGK-PuroR-loxP donor plasmid and 3 µg of PX458-LNCSOX17-promoter (1:1 molar ratio).

Two days post transfection, cells were selected with final 2 µg/ml Puromycin-Dihydrochloride (Thermo Fisher Scientific, A1113803) for 14 days at 37°C, 5% CO2. For the derivation of isogenic reporter cell lines, single-cell derived colonies were manually picked and expanded. Differentiation into EN followed by qRT-PCR analysis was used to confirm clones that were activating the reporter.

### 15. Generation of CRISPRi hiPS cell line

ZIP13K2 hiPSCs (s. *hiPS cell culture*) were treated with Accutase containing final 10 µM Y-27632 for 15 min at 37°C, 5% CO_2_ to obtain a single cell suspension. Cell suspensions were counted and seeded at a density of 1-2 x 10^5^ cells /cm^2^ in mTeSR1 supplemented with final 10 µM Y-27632. Cells were pre-cultured for 16 h at 37°C, 5% CO_2_ prior to transfection.

The following day, cells were transfected using Lipofectamin Stem Transfection Reagent in fresh mTeSR1 supplemented with final 10 µM Y-27632 for 24 h at 37°C, 5% CO_2_. Transfection mixtures contained 2 µg of Super PiggyBac transposase expression vector (SBI, PB210PA-1) and 4 µg dCas9-KRAB-MeCP2 (*51*) (1:1 molar ratio). dCas9-KRAB-MeCP2 was a gift from Alejandro Chavez & George Church (Addgene plasmid # 110821; http://n2t.net/addgene:110821; RRID:Addgene_110821).

Two days post transfection, cells were selected with final 2 µg/ml Blasticidin-S-HCl (Thermo Fisher Scientific, A1113903) for 14 days at 37°C, 5% CO_2_. For the derivation of isogenic CRISPRi cell lines, single-cell derived colonies were manually picked and expanded. IF stainings for Cas9 confirmed homogenous dCas9-KRAB-MeCP2 expression in the selected clones (s. *Immunofluorescence staining* for detailed experimental procedure).

### 16. Production of lentiviral particles carrying sgRNAs

Lentiviral particles of specific sgRNA constructs have been produced in HEK-293T cells by co-transfection of 1:1:1 molar ratios pCMV-VSV-G plasmid (addgene, #8454 (*101*), 3,5µg), psPAX2 plasmid (addgene, #12260, 7µg) in combination with sgRNA specific variants of pU6-sgRNA EF1Alpha-puro-T2A-BFP (*52*) plasmid (addgene, #60955, 14µg). pCMV-VSV-G was a gift from Bob Weinberg (Addgene plasmid # 8454; http://n2t.net/addgene:8454; RRID: Addgene_8454). Prior to transfection, HEK-293T cells were grown on a 10 cm dish up to 70-80% confluency in HEK-media (KO-DMEM (Themro Fisher Scientific, 10829018), 10% fetal bovine serum (FBS, PAN Biotech, P30-2602), 1 x GlutaMAX Supplement, 100 U/ml Penicillin-Streptomycin (Thermo Fisher Scientific, 15140122) and final 1 x, 5,5 µM ß-Mercaptoethanol (Thermo Fisher Scientific, 21985023)). For each sgRNA construct, plasmid DNA mixtures and 50 µl of LipoD293 transfection reagent (SignaGen Laboratories, SL100668) were mixed in 250 µl KO-DMEM at RT. After pipette mixing, transfection particles were incubated at RT for 15 min. Each sgRNA-specific mixture was added drop-wise onto HEK-293T cultures in 10 ml HEK-media and incubated for 16h at 37°C, 5% CO_2_. Cell culture media was exchanged by10 ml fresh HEK-media the next day and culture supernatants (S/N) of the two subsequent days were then filtered (0,22 µm), collected and stored at 4°C. After the second harvesting day, S/N were supplemented with 1 x PEG-it virus precipitation solution (SBI, LV810A-1) for 24 h at 4°C. Viral particles were finally precipitated by centrifugation at 3234 x g, 4°C. Viral precipitates were resuspended in 200 µl mTeSR1 and either frozen at −80°C or immediately used for lentiviral transduction of CRISPRi hiPSCs. The entire lentivirus preparation and storage was carried out under S2-safety conditions and pre-cautions.

### 17. Lentiviral transduction of CRISPRi hiPSCs

Lentiviral particles were either thawed on ice (if frozen) or directly used fresh on the day of production. For hiPS cells transduction, clump-based hiPSCs splitting was performed (s. *hiPS cell culture* for detailed experimental procedure) and dissociated clumps were supplemented with 10 µM Y-27632, 10 µg/ml Polybrene infection reagent (MerckMillipore, TR-1003-G) and 100 µl lentiviral particles preparation. Cells were then plated and cultured for 16 h at 37°C, 5% CO_2_. The following day, cells were washed 10 times with DPBS and given fresh mTeSR1 supplemented with 10 µM Y-27632 for 24 h at 37°C, 5% CO_2_.

Successfully infected cells were then selected with 2 µg/ml Puromycin Dihydrochloride (Thermo Fisher Scientific, A1113803) for 14 days at 37°C, 5% CO_2_. CRISPRi cell line expressing sgRNAs (sgLNCSOX17 and sgCtrl), were grown as bulk cultures, and Tag-BFP was used as a proxy for sgRNA expression prior to differentiation into the respective endodermal derivate.

### 18. RNA isolation and cDNA synthesis

For RNA extraction, cells were lysed in 500 μl Qiazol from the miRNeasy Mini Kit (Quiagen, 217004), followed by vortexing. RNA was then extracted using the miRNeasy Mini Kit (Quiagen, 217004) and RNA concentration was measured. cDNA synthesis was performed using 1 μg total RNA for each sample using the RevertAid First Strand cDNA Synthesis Kit (Thermo Fisher Scientifc, K1622), following the manufacturer’s instructions Random hexamers have been used as primers for first strand cDNA synthesis.

### 19. Quantitative PCR (qPCR)

Quantitative PCR (qPCR) was carried out on a StepOnePlus 96-well or a QuantStudio 7 Flex 384-well Real-Time PCR System (Thermo Fisher Scientific) loading 20-25ng cDNA /well and using TaqMan Fast Advanced Master-Mix (Thermo Fisher Scirentific, 4444557) with TaqMan validated probes (Table 3) (Thermo Fisher Scientific) following the manufacturer’s instructions.

### 20. 5’/3’ RACE PCR experiments

5’/3’ rapid amplification of cDNA ends (RACE) PCR reactions where performed utilizing the 5′/3′ RACE Kit, 2^nd^ Generation (Sigma-Aldrich, 3353621001) according to the manufacturer’s instructions. Corresponding gene specific (SP) primers are listed in Table 3. RACE-PCR products were cloned into pJET1.2 backbone followed by bacterial transformation and sanger sequencing.

### 21. Extraction of polyA RNA for Nanopore sequencing

Isolation of poly(A)-enriched mRNA was performed using the Dynabeads mRNA DIRECT purification kit (Thermo Fisher Scientific, 61011) according to the manufacturer’s instruction with minor modifications. ZIP13K2-derived EN cells were washed once with DPBS and dissociated with Accutase for 15 min at 37°C, 5% CO_2_. Enzymatic reaction was quenched by adding mTeSR1 and cells were counted using the Countess II automated cell-counter. A total of 4 x 10^6^ viable cells were centrifuged for 5 min at 4°C, 300 x g. The supernatant was discarded and cells were washed with 1 ml of ice-cold DPBS and centrifuged as described above. The supernatant was completely removed and the cell pellet was carefully resuspended in 1,25 ml Lysis/Binding buffer. In order to reduce viscosity resulting from released genomic DNA, the samples were passed through a 21 gauge needle (Becton Dickinson, 304432) for five times and subsequently added to the pre-washed Oligo(dT)_25_ beads. Hybridization of the beads/mRNA complex was carried out for 10 min on a Mini Rotator (Grant-bio) and vials were placed on a DynaMag2 magnet (Thermo Fisher Scientific, 12321D) until the beads were fully immobilized. The DNA containing supernatant was removed and the beads were resuspended twice with 2 ml of Buffer A following a second wash step with two times 1 ml of Buffer B. Purified RNA was eluted with 10 µl of pre-heated Elution Buffer (10 mM Tris-HCl pH 7,5) for 5 min at 80°C and quantified with a Qubit Fluorometer (Thermo Fisher Scientific) using the RNA HS Assay Kit (Thermo Fisher Scientific, Q32852). Eluted RNA samples were immediately used for preparation of Nanopore sequencing libraries or kept at −80°C.

### 22. Preparation of Nanopore sequencing libraries

Preparation of RNA sequencing libraries was performed following the manufacturer’s instructions (ONT, SQK-PCS109) with minor modifications. Briefly, 50 ng of freshly prepared poly(A)-enriched mRNA was subjected to reverse transcription and strand-switching reaction. A total of four PCR reactions, each containing 5 µl of reverse transcribed cDNA, was used for the attachment of rapid primers (cPRM). Sufficient amplification of long cDNA molecules was enabled by setting the PCR extension time to 19 min and a total of 12 x cycles were used for amplification. Samples were treated with 1 µl of Exonuclease I (New England Biolabs, M0293S) and subsequently pooled for SPRI bead cleanup. Wash steps were performed using 80% ethanol solution and beads were eluted in 60 µl of 50°C pre-heated nuclease-free water. Samples were then incubated for additional 20 min at 50°C. Eluted DNA was combined with 5 µl adapter mix (AMX), 25 µl ligation buffer (LNB) from ONTs ligation sequencing kit (ONT, SQK-LSK109) and 10 µl of NEBNext Quick T4 DNA Ligase (New England Biolabs, E6056S). Ligation mix was incubated at RT for 30 min. Removal of short DNA fragments was achieved by adding 40 µl of Agencourt AMPure XP beads (Beckmann Coulter, A63881) combined with 2 wash steps with 250 µl of long fragment buffer (LFB) included in ONTs ligation sequencing kit. The final library was eluted with 13 µl elution buffer (EB) for 20 min at 48°C and DNA concentration was quantified using the Qubit dsDNA BR assay kit (Thermo Fisher Scientific, Q32850). A total of 400 ng was carefully mixed with 37,5 µl sequencing buffer (SQB), 25,5 µl of loading beads (LB) and loaded onto a primed MinION flow cell (ONT, R9.4.1 FLO-MIN106).

### 23. RNA sequencing

ZIP13K2 hiPSCs and their derived EN cultures were treated with Accutase for 15 min at 37°C, 5% CO_2_ to obtain a single cell suspension. Cells were then collected, washed with ice cold DPBS and centrifuged at 4°C, 300 x g for 5 min. Subsequently, 350 µl of RLT Plus buffer containing 1% β-mercaptoethanol (Thermo) was added to the cell pellets for cell lysis. After dissociation by trituration and vortexing, RNA was extracted using RNeasy Plus Micro Kit (Qiagen) and RNA concentration and quality was measured using the Agilent RNA 6000 Pico Kit (Agilent Technologies, 5067-1513) on an Agilent 2100 Bioanalyzer. All samples analyzed had a RINe value higher than 8,0, and were subsequently used for library preparation. mRNA libraries were prepared using KAPA Stranded RNA-Seq Kit (KapaBiosystem) according to the manufacturer’s instructions. 500 ng of total RNA was used for each sample to enter the library preparation protocol. For adapter ligation dual indexes were used (NEXTFLEX® Unique Dual Index Barcodes NOVA-514150) at a working concentration of 71nM (5 µl of 1 uM stock in each 70 µl ligation reaction). Quality and concentration of the obtained libraries were measured using Agilent High Sensitivity D5000 ScreenTape (Agilent-Technologies, 5067-5592) on an Agilent 4150 TapeStation. All libraries were sequenced using 100 bp paired-end sequencing (200 cycles kit) on a NovaSeq platform at a minimum of 25 million fragments /sample.

### 24. 4C sequencing

Triplicates of either undifferentiated ZIP13K2 or ZIP13K2-derived EN cultures were collected as described previously. ZIP13K2-derived EN cultures were further quenched with MACS-buffer (Final DPBS, 2 mM EDTA (ThermoFisher Scientific), 0,5% BSA (Sigma-Aldrich)) to obtain a single cell suspension. CXCR4^+^ cell populations, were enriched using MicroBead Kit (Miltenyi Biotec) following the manufacturer’s instructions. Pre- and post-MACS enriched cell fractions of differentiated cultures were measured for CXCR4-APC signal on the FACS Aria II (Beckton Dickinson) to confirm the cell population purity. Circularized Chromosome Conformation Capture (4C) library preparation of undifferentiated, or differentiated CXCR4^+^ enriched cell populations was performed according to the Weintraub A.S. et al. protocol (*102*). Briefly, NlaIII (New England Biolabs, R0125) was used as the primary cutter and DpnII (New England Biolabs, R0543) as a secondary cutter. Touchdown PCR on 4C libraries was performed using specific primer-pairs (s. primer list in Table 3) for the respective view-points. Illumina sequencing libraries were then prepared and sequenced using 150 paired-end sequencing (300 cycles kit) on a HiSeq4000 platform at a minimum of 10M fragments/ sample.

### 25. Capture Hi-C sequencing

cHi-C libraries were prepared from CRISPRi sgCtrl or sgLNCSOX17 EN cells. 5 x 10^6^ ZIP13K2-derived EN cells were harvested and washed with ice cold DPBS. Cell lysis, NlaIII (NEB, R0125) digestion and proximity-ligation was performed according to the Franke et al. protocol (*103*) with minor changes. Adaptors were added to DNA fragments and amplified according to Agilent Technologies instructions for Illumina sequencing. The library was hybridized to the custom-designed SureSelect probes (Agilent Technologies, 5190-4806 / 3253271) (s. probe list in Table 3) and indexed for sequencing of 200 M fragments /sample (100 bp paired-end) following the Agilent instructions. Capture Hi-C experiments were performed as biological duplicates.

### 26. SOX17 Chromatin Immunoprecipitation (ChIP) sequencing

ZIP13K2-derived EN cells (5 x 10^6^ /IP) were harvested and cross-linked in 1% formaldehyde (Thermo Fisher Scientific, 28908) in DPBS for 10 min at RT, followed by quenching with final 125 mM Glycine (Sigma-Aldrich, 50046) for 5 min at RT. Cross-linked cells were then centrifuged at 500 x g at 4°C and washed twice with ice cold DPBS. Cell lysis was performed by resuspending the pellet in 500 μl Cell Lysis Buffer (Final 10mM Tris-HCl, pH 8,0 (Sigma Aldrich, T2694); 85mM KCl (Sigma Aldrich, P9541); 0,5% NP40 (Sigma Aldrich, 56741); 1 x cOmplete, EDTA-free Protease Inhibitor Cocktail (Sigma Aldrich, 11873580001)) followed by 10 min incubation on ice. After the incubation, lysed cells were centrifuged at 2500 x g for 5 min at 4°C. Supernatant was carefully removed and the extracted nuclei were then resuspended in 230 μl Nuclei Lysis Buffer (Final 10mM Tris-HCl, pH 7,5 (Sigma Aldrich, T2319)); 1% NP40; 0,5% sodium deoxycholate (Sigma Aldrich, D6750); 0,1% SDS (Thermo Fisher Scientific, AM9820); 1 x cOmplete, EDTA-free Protease Inhibitor Cocktail). Following 10 min incubation on ice, each 260 μl sample was split into two microTUBEs (Covaris, 520045) and chromatin was sonicated using a Covaris E220 Evolution with the following settings: Temperature → 4°C; Peak power → 140; Duty factor → 5,0; Cycles/Burst → 200; Duration → 750 sec. After sonication, sheared chromatin (ranging from 200-600bp) was transferred in a new 1,5 ml tube and centrifuged at max speed for 10 min at 4°C. Supernatant was then transferred into a new tube and volume was increased to 1 ml /sample with ChIP Dilution Buffer (Final 16,7mM Tris-HCl, pH 8,0; 1,2mM EDTA (Sigma Aldrich, 03690)); 167mM NaCl (Sigma Aldrich); 1,1% Triton-X (Sigma Aldrich); 0,01% SDS; 1 x Protease Inhibitor). 50μl (5%) was then transferred into a new tube and frozen at −20°C as INPUT. 1μg of SOX17 antibody /10^6^ initial cells was added to the 950 μl left, and immunoprecipitation was carried out at 4°C o/n on a rotator (s. Table 3). The next day, 50μl of Dynabeads Protein G (Thermo Fisher Scientific, 10004D) /IP were washed twice with ice cold ChIP Dilution Buffer and then added to each IPs. IP/bead mixes were incubated for 4 hours at 4°C on a rotor. Next, bead/chromatin complexes were washed twice with Low Salt Wash Buffer at 4°C (Final 20 mM Tris-HCl, pH 8,0; 2 mM EDTA; 150 mM NaCl (Sigma-Aldrich, S6546); 1% Triton-X; 0,1% SDS), twice with High Salt Wash Buffer at 4°C (Final 20 mM Tris-HCl, pH 8,0; 2 mM EDTA; 500 mM NaCl; 1% Triton-X; 0,1% SDS), twice with LiCl Wash Buffer at 4°C (Final 10 mM Tris-HCl, pH 8,0; 1mM EDTA; 250mM LiCl (Sigma Aldrich, L9650); 1% sodium deoxycholate (Sigma Aldrich); 1% NP40), twice with TE pH 8,0 (Sigma Aldrich, 8890) at room temperature and finally eluted twice in 50 μl freshly prepared ChIP Elution Buffer (Final 0,5% SDS; 100 mM NaHCO3 (Sigma Aldrich, S5761)) at 65°C for 15 min (total 100 μl final eluent). Thawed INPUTS and eluted IPs were next reverse cross-linked at 65°C o/n after the addition of 16 ul freshly prepared Reverse Crosslinking Salt Mixture (Final 250 mM Tris-HCl, pH 6,5 (Sigma Aldrich, 20-160); 62,5 mM EDTA; 1,25M NaCl; 5 mg/ml Proteinase K (Thermo Fisher Scientific, AM2548)). The following day phenol:chloroform (Thermo Fisher Scientific, 15593031) extraction followed by precipitation was performed to isolate DNA. IPs and INPUTS were then quantified and NGS libraries were prepared using NEBNext Ultra II DNA Library Prep Kit for Illumina (New England Biolabs, #E7645) following the manufacturer’s instructions. Library quality and size distribution was verified using a TapeStation D5000 HS kit (Agilent Technologies, 5067-5592). Samples were sequenced with a coverage of 50 M paired end reads (2 x 100 bp) /sample on a NovaSeq (Illumina).

### 27. GATA4/GATA6 Chromatin Immunoprecipitation (ChIP) sequencing

GATA4/6 ChIPs were perfored in duplicates as previously described (*104*). Briefly, approximately 5×106 cells were used for each IP. Cells were cross-linked with 1% formaldehyde for 10 minutes followed by quenching with 125 mM glycine for 4-5 minutes at room temperature. The cell pellet was lysed in cell lysis buffer (20 mM Tris-HCl pH 8, 85 mM KCl, 0.5% NP-40) supplemented with 1X protease inhibitors (Roche, 11836170001) on ice for 20 minutes then spun at 5000 rpm for 10 minutes. The nuclear pellet was resuspended in sonication buffer (10 mM Tris pH 7.5, 1% NP-40, 0.5% sodium deoxycholate, 0.1% SDS, and 1X protease inhibitors) and incubated for 10 minutes at 4°C. In order to achieve a 200-700 bp DNA fragmentation range, nuclei were sonicated using a Bronson sonifier (model 250) with the following conditions: amplitude = 15%, time interval = 3min (total of 8-12 minutes) and pulse ON/OFF = 0.7 s/1.3 s. Chromatin was pre-cleared with Dynabeads Protein A (Invitrogen, 10002D) for 1 hour and incubated with antibody on a rotating wheel overnight at 4°C. On the following day, 30-40 μl of Dynabeads Protein A was added to chromatin for 2-3 hours. The captured immuno-complexes were washed as follows – 1x in low-salt buffer, 1x in high-salt buffer, 1x in LiCl salt buffer, and 1x in TE. The immuno-complexes were eluted in ChIP-DNA elution buffer (10 mM Tris-HCl pH 8, 100 mM NaCl, 20 mM EDTA, and 1% SDS) for 20 minutes. The eluted ChIP-DNA was reverse cross-linked overnight at 65°C, followed by proteinase K (Thermo, 25530049) treatment, RNase A (Thermo, ENO531) treatment, and Phenol:Chloroform:Isoamyl alcohol extraction. The Illumina library construction steps were carried out with 5-10 ng of purified DNA. During library construction, purification was performed after every step using QIAquick PCR purification kit (QIAGEN, 28104) or QIAquick gel extraction kit (QIAGEN, 28706). The library reaction steps were as follows: end-repair, 3′ end A-base addition, adaptor ligation, and PCR amplification. The amplified libraries were size-selected for 200-450 bp on a 2% agarose E-gel (Thermo, G402002) and sequenced (single-end, 75) on a NextSeq500 or Hi-Seq2000 platform.

### 28. H3K9me3 Chromatin Immunoprecipitation (ChIP) qPCR

ZIP13K2-derived EN cells (2 x 10^6^ /IP) were harvested, cross-linked, washed, lysed and sonicated as described previously (s. *SOX17 ChIP sequencing*). ChIP for H3K9me3 was performed in triplicates utilizing the High-Sensitivity ChIP Kit (abcam, ab185913) in combination with the ChIP-grade H3K9me3 antibody (ab8898, abcam) according to the manufacturer’s instructions with slight modifications. Instead of DNA column purification, phenol:chloroform extraction followed by precipitation was performed to isolate DNA (s. *SOX17 ChIP sequencing*). Precipitated DNA was dissolved in 200 µl H_2_O.

qPCR reactions were set up utilizing the 2 x PowerUp SYBR Green Master Mix (Thermo Fisher Scientific, A25777) containing final 250 nM forward /reverse primer (s. Table 3). All samples have been measured in technical triplicates using 4 µl diluted input or IP sample from above /reaction/replicate. qPCRs were set-up on 96-well plates (Thermo Fisher Scientific, N8010560), spun down for 1 min at 2500 x g, RT and ran on a StepOnePlus 96-well Real-Time PCR System (Thermo Fisher Scientific).

### 29. *LNCSOX17* RNA-pulldown followed by Mass Spectrometry

RNA-pulldown protocol to discover *LNCSOX17* protein interaction partners has been performed combining (*94, 105*) protocols with some modifications. ZIP13K2-derived EN cells (60 x 10^6^) were harvested and cross-linked in 1% formaldehyde (Thermo Fisher Scientific, 28908) in DPBS for 5 min at RT, followed by quenching with final 125 mM Glycine (Sigma-Aldrich, 50046) for 5 min at RT. Cross-linked cells were then centrifuged at 500 x g at 4°C and washed three times with ice cold DPBS. Cells are then resuspended in 10ml Sucrose/Glycerol buffer (1:1) (*Sucrose Buffer*: 0.3M Sucrose; 1% Triton-X (Sigma Aldrich); 10mM HEPES (Thermo Fisher Scientific, 31330038); 100mM KOAc; 0,1 mM EGTA (Sigma Aldrich); 0,5mM Spermidine; 0,15mM Spermine; 1mM DTT; 1X proteinase inhibitor (Roche, 11836170001); 10U/ml SUPER-asIN (Thermo Fisher Scientific, AM2694)) (*Glycerol Buffer*: 25% Glycerol; 10mM HEPES; 100mM KOAc; 0,1mM EGTA; 1mM EDTA (Sigma Aldrich, 03690); 0,5mM Spermidine; 0,15mM Spermine; 1mM DTT; 1X proteinase inhibitor; 10U/ml SUPER-asIN) and dounced 20 times in a glass tight pestle (Sigma Aldrich, D9938-1SET). After douncing, lysed cells are incubated for 10 min on ice inside the pestle. Cells are then transferred on a cushion of 10 ml Glycerol Buffer in a 50 ml falcon tube and centrifuged at 1000 x g for 15 min at 4°C to recover nuclei. Supernatant is discarded by pipetting first, and residual volume is decanted on a clean paper towel. Extracted nuclei are then resuspended in 5 ml 3% formaldehyde and fixed again for 30 min at RT, followed by three DPBS washes. Next, nuclei are resuspended in 5 ml Nuclei Extraction Buffer (Final 50mM HEPES, pH 7,5; 250mM NaCl; 0,1% sodium deoxycholate (Sigma Aldrich, D6750); 0,1mM EGTA; 0,5% N-lauroylsarcosine; 5mM DTT; 100 U/ml SUPER-asIN) and incubated for 10 min on ice. Nuclei are then centrifuged at 400 x g for 5 min at 4°C, and resuspended in 530 ul Nuclei Resuspension Buffer (Final 50mM HEPES, pH 7,5; 75mM NaCl; 0,1% sodium deoxycholate; 0,1mM EGTA; 0,5% N-lauroylsarcosine; 5mM DTT; 100 U/ml SUPER-asIN) and sonicated using a Covaris E220 Evolution with the following settings: Temperature → 4°C; Peak power → 140; Duty factor → 5,0; Cycles/Burst → 200; Duration → 15 mins. After sonication, sheared chromatin is split into 3 samples (Even/Odd/LacZ, 120ul each) and incubated with the corresponding biotinylated probes set (36 pmols of probes are added; see Table 3 for probes sequences) together with 240ul Hybridization Buffer (Final 33mM HEPES, pH 7,5; 808mM NaCl; 0,33% SDS; 5mM EDTA; 0,17% N-lauroylsarcosine; 2,5mM DTT; 5X Denhardt’s solution; 1X proteinase inhibitor; 100 U/ml SUPER-asIN) overnight at RT on a rotor. 5% sonicated sample was frozen as INPUT. The next day, 240μl of MyOne Streptavidin C1 beads (Thermo Fisher Scientific, 65001) were added to each pulldown after washing and resuspension in Hybridization Buffer, and incubated for 3 h at RT on a rotor. Next, bead complexes were washed once with Wash Buffer 1 (Final 30mM HEPES, pH 7,5; 1,5 mM EDTA; 240mM NaCl; 0,75% N-lauroylsarcosine; 0,65% SDS; 0,7mM EGTA; 2M Urea), four times with Wash Buffer 2 (Final 10mM HEPES, pH 7,5; 2 mM EDTA; 240mM NaCl; 0,1% N-lauroylsarcosine; 0,2% SDS; 1mM EGTA) and once with RNase H elution Buffer (Final 50mM HEPES, pH 7,5; 1,5 mM EDTA; 75mM NaCl; 0,125% N-lauroylsarcosine; 0,5% Triton-X; 10mM DTT; 0,5M Urea). In this last step, 10% of the beads from each pulldown is transferred to a new tube for RNA isolation. The remaining 90% (protein sample fraction) is eluted in RNase H elution Buffer containing 10% RNase H, 10% RNase A and 10% DNase for 30 min at RT. The RNase fraction is de-crosslinked together with the INPUT samples with Proteinase K (Thermo Fisher Scientific, AM2548) treatment and RNA is extracted following Trizol purification. RNA and INPUT samples were reverse transcribed and used for qPCR to validate LNCSOX17 enrichment. Protein samples were run on a NuPAGE 4-12%, Bis-Tris, 1,0 mm, Mini Protein Gel, Silver stained using SilverQuest (Thermo Fisher Scientific; LC6070) following manufacturer instructions. The mass spectrometry compatible SilverQuest Silver Staining Kit was used for de-staining. Gel pieces were then washed twice with 300 µL of 25mM ammonium bicarbonate in 50% acetonitrile, shaking at 500 rpm for 10 min, followed by centrifugation at 16,000 x g for 30 sec. Gel pieces were completely dried in a vacuum concentrator. In-gel digestion with trypsin and extraction of peptides was done as previously described (*106*). Dried peptides were reconstituted in 5% acetonitrile and 2% formic acid in water, briefly vortexed, and sonicated in a water bath for 30 sec before injection to nano-LC-MS. LC-MS/MS was carried out by nanoflow reverse-phase liquid chromatography (Dionex Ultimate 3000, Thermo Scientific) coupled online to a Q-Exactive HF Orbitrap mass spectrometer (Thermo Scientific), as reported previously (*107*). Briefly, the LC separation was performed using a PicoFrit analytical column (75μm ID × 50cm long, 15µm Tip ID; New Objectives, Woburn, MA) in-house packed with 3-µm C18 resin (Reprosil-AQ Pur, Dr. Maisch, Ammerbuch, Germany). Peptides were eluted using a gradient from 3.8 to 38% solvent B in solvent A over 120 min at a 266nL per minute flow rate. Solvent A was 0.1% formic acid and solvent B was 79.9% acetonitrile, 20% H_2_O, and 0.1% formic acid. For the IP samples, a one-hour gradient was used. Nanoelectrospray was generated by applying 3.5kV. A cycle of one full Fourier transformation scan mass spectrum (300−1750m/z, resolution of 60,000 at m/z 200, automatic gain control (AGC) target 1×10^6^) was followed by 12 data-dependent MS/MS scans (resolution of 30,000, AGC target 5×10^5^) with a normalized collision energy of 25eV.

### 30. HNRNPU RNA Immunoprecipitation (RIP) followed by qRT-PCR or Western Blot

ZIP13K2-derived EN cells (10 x 10^6^) were harvested and cross-linked according to the manufacturer’s instructions in 0,3% formaldehyde in DPBS for 10 min at RT, followed by quenching with final 1 x Glycine solution for 5 min at RT utilizing the Magna Nuclear RIP (Cross-Linked) Nuclear RNA-Binding Protein Immunoprecipitation Kit (Merck millipore, 17-10520). Cross-linked cells were then centrifuged at 800 x g at 4°C and washed three times with ice cold DPBS. Supernatant free cell pellets were conducted to cell lysis according to the Kit manufacturer’s instructions. Sonication has been performed in Kit provided RIP Cross-linked Lysis Buffer using the Covaris E220 Evolution with the following settings: Temperature → 4°C; Peak power → 140; Duty factor → 5,0; Cycles/Burst → 200; Duration → 6 mins. to obtain a DNA smear of 200-1000 bp. Sonicated lysates were centrifuged at 1000 x g for 10 min at 4°C and supernatants aliquoted and stored at −80°C. DNase I treatment following Immunoprecipitation has been performed according to the Kit manufacturer’s instructions, combining lysates corresponding to 10^6^ cells with 5 µg antibody per sample (s. Table 3 for antibodies). After DNase I treatment 10% input material for qRT-PCR has been kept and stored at −80°C. Initial supernatants (unbound fraction w/o beads) after o/n immunoprecipitation and 10% material of the last wash step (IP including beads) has been kept for Western Blot and stored at −20°C. Inputs and IP were further conducted to reverse crosslinking and RNA purification according to the Kit manufacturer’s instructions. cDNA synthesis has been carried out as mentioned earlier (s. *RNA isolation and cDNA synthesis*).

Quantitative PCR (qPCR) reactions were set up utilizing the 2 x PowerUp SYBR Green Master Mix (Thermo Fisher Scientific, A25777) containing final 250 nM forward /reverse primer (s. Table 3 for primer & oligos) and 20-25ng cDNA /well. Reactions were set up in 384-well plates (Thermo Fisher Scientific, AB2384B) following centrifugation for 2 min at 2500 x g, RT. Reactions were carried out on a QuantStudio 7 Flex 384-well Real-Time PCR System (Thermo Fisher Scientific).

Western Blot samples of unbound fractions and IP were boiled in final 1 x Laemmli Buffer (BioRad, 1610747) containing 10% 2-Mercaptoethanol (M6250, Sigma-Aldrich) for 10 min at 95°C, followed by cooling on ice for 5 min. Western Blots have finally been carried out as described below (s. *Western Blot*) utilizing respective antibody dilutions (s. Table 3 for antibodies).

### 31. Immunofluorescence staining

For immunofluorescent stainings, cells were grown in Ibidi 8-well glass-bottom plates (Ibidi, 80827) (initial seeding, 10^4^ cells /well). On the day of analysis, cells were washed twice with DPBS and then fixed in 4% Paraformaldehyde (PFA) solution (Sigma-Aldrich, P6148-500G) for 30 min at 4°C, and then washed three more times with DPBS. Subsequently, cells were permeabilized for 30 min in DPBS-T solution (Final 0,5% Triton-X (Sigma-Aldrich, T8787-50 ML) in DPBS) and blocked for 30 min in Blocking solution (Final 10% fetal bovine serum in DPBS-T) at RT. Primary antibody incubation was performed in blocking solution for 1 h and 45 min at RT, after which cells were washed three times with Blocking solution. After the last washing step, samples were incubated with secondary antibodies diluted in Blocking solution for 30 min at RT. Afterwards, cells were washed three times with DPBS-T. The last DPBS-T washing step after secondary antibody incubation contained 0,02% DAPI (Roche Diagnostics, 10236276001). DAPI was incubated for 10 min at RT and washed off once with DPBS. All primary and secondary antibodies and their working concentrations are listed in Table 3.

### 32. Cell clearing

Prior to imaging, cells were cleared with RIMS (Refractive Index Matching Solution) in order to increase light penetrability. To this end, samples were first washed three times with 0,1 M phosphate buffer (0,025 M NaH_2_PO_4_, 0,075 M Na_2_HPO_4_, pH 7,4). Clearing was then performed by incubation in RIMS solution (133% w/v Histodenz (Sigma-Aldrich, D2158) in 0,02 M phosphate buffer) at 4°C o/n.

### 33. Immunofluorescence imaging

Cells stained with antibodies were imaged with the Zeiss Celldiscoverer7 (wide-field), Zeiss LSM880 (laser-scanning microscope with Airyscan), Zeiss Observer (wide-field) or Nikon Eclipse TS2 (bench-top microscope) with appropriate filters for DAPI, Alexa Fluor 488, Alexa Fluor 568, Alexa Fluor 647, and combinations thereof.

### 34. Quantitative fluorescence microscopy

For each staining tested, a total of 49 individual positions were acquired in 3 fluorescence channels/replicate /well, with a 20 x /NA=0,95 objective, an afocal magnification changer 1 x, 3 x 3 camera binning, a consequential pixel size of 0,46 µm^2^, and in constant focus stabilization mode. Analysis was then performed using the Image Analysis module running in ZEN 3.2. On average 6928 single cells were analyzed per replicate. Cells were identified on smoothed nuclear counterstaining (DAPI) using fixed intensity thresholds, nearby objects were separated by mild water shedding. The consequential primary objects were filtered (area 45-175 µm^2^) and expanded by 8 pixels (=5,44µm^2^); the consecutive ring, surrogated a cytoplasm compartment. Fluorescence intensities (mean and standard deviation) were quantified for each nucleus and expanded object, depending on the staining pattern profiled.

### 35. Single molecule RNA fluorescent in situ hybridization

For single molecule RNA fluorescent in situ hybridization (smRNA-FISH), cells were grown in Ibidi 8-well glass-bottom plates (Ibidi 80827) (initial seeding, 10^4^ cells /well). On the day of analysis, cells were washed twice with DPBS, fixed in 4% PFA for 10 min at RT, and washed again twice with DPBS. Cells were then incubated in 70% ethanol at 4°C for at least 1 h and then washed with 1 ml of Wash Buffer A (LGC Biosearch Technologies) at room temperature for 5 min. Cells were subsequently hybridized with 100 μl of Hybridization Buffer (LGC Biosearch Technologies) containing the smRNA-FISH probes at a 1:100 dilution in a humid chamber at 37°C o/n (not more than 16 h). The next day, cells were washed with 1 ml of Wash Buffer A at 37°C for 30 min and stained with Wash Buffer A containing 10 μg/ml Hoechst 33342 at 37°C for 30 min. Cells were then washed with 1 ml of Wash Buffer B (LGC Biosearch Technologies) at RT for 5 min, mounted with ProLong Gold (Thermo, P10144), and left to curate at 4°C o/n before proceeding to image acquisition. Oligonucleotides probes were designed with the Stellaris smRNA-FISH probe designer (LGC Biosearch Technologies, version 4.2), labeled with Quasar 570 and produced by LGC Biosearch Technologies. smRNA-FISH probes sequences are listed in Table 3.

### 36. smRNA-FISH imaging

Image acquisition was performed using a DeltaVision Elite widefield microscope with an Olympus UPlanSApo 100 x /1,40-numerical aperture oil objective lens and a PCO Edge sCMOS camera. Z-stacks of 200 nm step size capturing the entire cell were acquired. Images were deconvolved with the built-in DeltaVision SoftWoRx Imaging software and maximum intensity projections were created using ImageJ (*108*) and Fiji (*109*).

### 37. Staining for FACS analysis

Undifferentiated or differentiated ZIP13K2 cultures were treated with Accutase for 15 min, 37°C, 5% CO_2_ to obtain a single cell suspension. To quench the dissociation reaction and to wash the cells, FACS-buffer was added (Final DPBS, 5 mM EDTA (ThermoFisher Scientific, 15575020), 10% Fetal bovine serum (FBS, PAN Biotech, P30-2602)). Next, cells were spun down at 300 x g, 5 min at 4°C. Cells were then resuspended in FACS-buffer containing surface marker antibodies (s. Table 3) and incubated for 15 min at 4°C in the dark. For extracellular stainings (ECS) only, cells were further washed once with FACS-buffer and spun down at 300 x g before FACS analysis was performed. If additional intracellular stainings (ECS+ICS) were performed, cells were washed once with FACS-buffer, supernatants were removed and cells fixed according to the manufacturer’s instructions utilizing the True-Nuclear™ Transcription Factor Buffer Set (Biolegend, 424401). Intracellular staining was performed according to manufacturer’s instructions before FACS analysis was carried out. ICS antibody dilutions are listed in Table 3. FACS analysis was performed on the FACSCelesta Flow Cytometer (Beckton Dickinson). Raw data were analyzed using FlowJo (LLC) V10.6.2.

### 38. Western Blot

Undifferentiated or differentiated ZIP13K2 cultures were treated with Accutase for 15 min, 37°C, 5% CO_2_ to obtain a single suspension. Single cell suspensions were washed once with ice cold DPBS and spun down at 300 x g, 5 min at 4°C. Supernatants were removed and cell lysates generated by treatment for 30 minutes on ice with RIPA buffer (Thermo Fisher Scientific, 89900) supplemented with 1 x HALT protease inhibitor (Thermo Fisher Scientific, 87786). Lysates were spun down at 12000 x g, 10 min at 4°C and supernatants quantified for protein content using the Pierce BCA Protein Assay Kit (Thermo Fisher Scientific, 23227) according to the manufacturer’s instructions.

For Western Blot, 20 µg total protein extract per sample were boiled in final 1 x Laemmli Buffer (BioRad, 1610747) containing 10% 2-Mercaptoethanol (M6250, Sigma-Aldrich) for 10 min at 95°C,followed by cooling on ice for 5 min. Samples were then loaded on a NuPAGE 4-12%, Bis-Tris, 1,0 mm, Mini Protein Gel (Thermo Fisher Scientific, NP0322BOX) and ran at 200 V for 30 min in 1 x NuPAGE MOPS SDS Running Buffer (Thermo Fisher Scientific, NP0001) containing 1:400 NuPAGE Antioxidant (Thermo Fisher Scientific, NP0005). Protein transfer has been performed utilizing the iBlot 2 Starter Kit, PVDF (Thermo Fisher Scientific, IB21002S) following the manufacturer’s instructions for the P0 program.

PVDF membranes containing transferred proteins were incubated in blocking buffer (1 x TBS-T (Thermo Fisher Scientific, 28360), 5% Blotting-Grade Blocker (BioRad, 1706404)) for 1 h at RT. Incubation with primary antibody dilution (s. Table 3) was performed in blocking buffer at 4°C overnight. The following day, membranes were washed three times 10 min at RT with 1 x TBS-T and incubated for 2 h at RT in secondary antibody dilution in blocking buffer (Table 3). Next, membranes were washed three times for 10 min at RT with 1 x TBS-T and developed using the SuperSignal West Dura Extended Duration Substrate (Thermo Fisher Scientific, 34075) according to the manufacturer’s instructions on the BioRad ChemiDoc XRS+ imaging system.

## Computational analysis

Command-line processing of BAM, BED and bigwig files was done using SAMtools (v1.10) (*110*), BEDtools (v2.25.0) (*111*) and UCSCtools (v4) (*112*). If not stated otherwise: All statistics and plots are generated using R version 3.6.0 and 3.6.1. In all boxplots, the centerline is median; boxes, first and third quartiles; whiskers, 1.5 x inter-quartile range; data beyond the end of the whiskers are displayed as points.

### 1. Human vs. mouse *LNCSOX17* conservation analysis

Local alignment was performed with EMBOSS Water (*113*). Visualizations were created with Matplotlib (*114*). Alignment sequences were read into python using the Biopython library (*115*). The full sequence of the human *LNCSOX17* locus was aligned to the full sequence of the mouse *Lncsox17* locus using Water. Aligned subsequences of 20 base pairs or more in length, including substitutions but excluding indels were used to calculate conservation. Additionally, individual exons and the enhancer sequence were also aligned with Water. Conserved stretches were connected from the human sequence box to the mouse sequence box and visualized as lines.

### 2. 4Cseq data analysis

The raw sequencing reads were trimmed by using cutadapt (*116*) (--discard-untrimmed -e 0.05 –m 25) to remove primer sequences and restriction enzyme sequences. The reads not matching those sequences, were removed from further analysis. The remaining reads were then mapped to the reference sequences GRCh37/hg19 by bowtie2 (*117*) (default parameters). An iterative mapping procedure was performed. Specifically, the full-length reads were first mapped to the genome. The unmapped reads were then cut by 5-nt from the 3-prime end each time until they were successfully mapped to the genome or until they were shorter than 25 bp. The final mapped reads were assigned to valid fragments. The fragment counts were then normalized by RPM (reads per million) and smoothed by averaging the counts of the closest 5 fragments.

### 3. Coding potential calculation

Whole genome multiple species alignments of 46 vertebrate species with human (assembly hg19, October 2009) as a reference have been retrieved from the UCSC genome browser (*118*). Human lincRNA annotation was obtained from Gencode (*119*) (gencode.v33lift37.long_noncoding_RNAs.gtf, December 2019). All ORFs in each transcript were identified and the corresponding multiple species alignment was scored by the omega method of PhyloCSF (*45*) (Figure 2C, left panel) shows 95% (2.5-97.5percentile) of the 271,572 sORFs from the (*118*) analyzed human lincRNAs. The *SOX17* CDS and all identified sORFs in *LNCSOX17* were scored by omega phyloCSF as shown in Figure 2C, right panel.

### 4. RNA-seq

All RNAseq samples were pre-processed using cutadapt (*116*) to remove adapter and trim low quality bases. Reads were subsequently aligned against the human reference genome hg19 using STAR (*120*) (parameter: outSAMtype BAM SortedByCoordinate --outSAMattributes Standard -- outSAMstrandField intronMotif --outSAMunmapped Within --quantMode GeneCounts). Finally, Stringtie (*121*) was used for calculation of strand-specific TPMs.

### 5. Capture Hi-C

Raw sequence reads of capture Hi-C (cHi-C) were mapped to the hg19 version of the human genome using BWA (v0.7.17-r1188) (*122*) with parameters (mem -A 1 -B 4 -E 50 -L 0). Mapped reads were further processed by HiCExplorer (v3.6) (*123*) to remove duplicated reads and reads from dangling ends, self-circle, self-ligation and same fragments. The replicates were merged to construct contact matrices of 1 kb resolution. Normalization was performed to ensure that all samples have the same number of total contacts, followed by KR correction. The relative contact difference between two cHi-C maps was calculated by subtracting one from the other using the corrected matrices.

### 6. SOX17 Chromatin Immunoprecipitation

The ChIP-seq sequencing data as well as the control input sequencing were aligned to the human reference genome (hg19) using BWA mem (*124*) using the default parameter. GATK (*125*) was used to obtain alignment metrics and remove duplicates. Peaks were called using the MACS2 (2.1.2_dev) (*126*) peakcall function using default parameters. After validation of replicate comparability and quality, replicates were merged on read level and reprocessed together with input samples. Background subtracted coverage files were obtained using MACS2 bdgcomp with -m FE.

### 7. GATA4/6 Chromatin Immunoprecipitation

The ChIP-seq sequencing data as well as the Fastqs for GATA4/6 ChiP-seq experiments were processed using the ENCODE ChIP-seq pipeline version 1.6.1 (https://github.com/ENCODE-DCC/chip-seq-pipeline2) using default settings with the hg19 genome. Standard ENCODE ChIP-seq reference files were used as found in https://storage.googleapis.com/encode-pipeline-genome-data/genome_tsv/v1/hg19_caper.tsv. Pooled fold-change bigWigs were used.

### 8. Single-cell RNAseq pipeline

Publicly available single-cell RNAseq raw data of already filtered 1195 cells from a gastrulating human embryo (*44*) was downloaded from ArrayExpress (*127*) under accession code E-MTAB-9388. The GENCODE (*128*) human transcriptome (GRCh37.p13) and its annotation were downloaded and added with the *LNCSOX17* entry. After building the transcriptome index, the transcripts abundance was quantified via Salmon v1.6.0 (*129*) in quasi-mapping-based mode using the –seqBias and the – gcBias flags. Data was loaded as a scanpy v1.4.4 (*130*) object, reproducing clustering as reported by Tyser, R. C. v. et al. (*44*). The resulting clusters were visualized via the scanpy UMAP representation in two dimensions, using default parameters (tl.umap). UMAPs are displayed in Figure supplement 2E (upper panel).

### 9. Bulk measurements from scRNAseq pipeline

To measure LNCSOX17 read counts in endoderm cells fastq files were combined in one bulk raw file. The file went through a bulk RNAseq pipeline comprising a pre-alignment quality control via fastQC v0.11.9, adaptor and low-quality bases trimming using cutadapt (*116*), post-QC and reads alignment against the human genome (GRCh37.p13) by means of STAR (*120*) (parameters: -- outSAMtype BAM SortedByCoordinate, --chimSegmentMin 20, --outSAMstrandField intronMotif, --quantMode GeneCounts). Finally, the BAM file was visualized using the Integrative Genomic Viewer (IGV) (*131*). IGV tracks are displayed in Figure supplement 2E (lower panel).

### 10. Oxford Nanopore RNA analysis

All Oxford Nanopore Technologies derived runs were processed using the Nanopype pipeline (v1.1.0) (*132*). The basecaller Guppy (v4.0.11) was used with the r9.4.1 high-accuracy configuration. Quality filtering was disabled for any base calling. Base-called reads were aligned against the human reference genome hg19 using minimap2 (v2.10) (*133*) with the Oxford Nanopore Technologies parameter preset for spliced alignments (-ax splice -uf -k14). Only unique alignments (-F 2304) are reported.

### 11. Oxford Nanopore RNA split-read analysis

Nanopore post processed split read data (*s. Oxford Nanopore RNA analysis*) from wild type endoderm mRNA (*s. Extraction of polyA RNA for Nanopore sequencing; s. Preparation of Nanopore sequencing libraries*) were extracted from the junctions-track of BAM files visualized using the Integrative Genomic Viewer (IGV) (*131*) utilizing the coordinates hg19, chr8:55115873-55141447. Split reads between hg19, chr8:55140801 (5’-sequence of Exon 1, *s. 5’/3’ RACE PCR experiments*) and hg19, chr8: 55125601 (3’-sequence of Exon 3, *s. 5’/3’ RACE PCR experiments*) were accounted for isoform Ex1+2 (s. Figure supplement 1C). Full isoform Ex1+2 sequence (∼2,8kb long) can be found in Table 1.

Split reads between hg19, chr8:55140801 (5’-sequence of Exon 1, *s. 5’/3’ RACE PCR experiments*) and hg19, chr8:55123254 (3’-sequence of Exon 3, *s. 5’/3’ RACE PCR experiments*) were accounted for isoform Ex1+3 (s. Figure supplement 1C). All other reads were accounted as “sloppy spliced” reads and together with both isoforms calculated in relative terms (s. Figure supplement 1C). Full isoform Ex1+3 sequence (∼3kb long) can be found in Table 1. Summary of the relative isoform quantification is displayed in Figure 2F.

### 12. RNA secondary structures prediction

Secondary structures for both isoforms of LNCSOX17 have been computed using the RNAfold tool in the Vienna RNA Websuite package (*134, 135*). The minimum free energy (MFE) structures have been calculated based on the partition function and base pairing probability matrix.

### 13. Mass spectrometry analysis and ranking of *LNCSOX17* protein partners

Raw MS data were processed with MaxQuant software (v 1.6.10.43) and searched against the human proteome database UniProtKB with 75,074 entries, released in May 2020. Parameters of MaxQuant database searching were a false discovery rate (FDR) of 0.01 for proteins and peptides, cysteine carbamidomethylation was set as fixed modification, while N-terminal acetylation and methionine oxidation were set as variable modifications. Protein abundance in each of the three samples has been quantified by calculating Label free quantitation (LFQ) values for each detected protein. Protein targets have then been ranked based on the Log_2_[(Even_LFQ_+Odd_LFQ_)/2)/LacZ_LFQ_] extrapolated values.

#### Plotting

Plots were generated with GraphPad Prism 8, R 3.6.0 and R 3.6.1.

## DECLARATIONS

### Ethics approval and consent to participate

Not applicable.

### Consent for publication

Not applicable.

### Availability of data and materials

All data presented in this study are available in the main text, methods or tables. Sequencing data have been deposited in the Gene Expression Omnibus (GEO) under accession code GSE178990.

Codes used to perform the analysis in this study are available upon request.

## Author contributions

A.L., A.B. and A.M. conceived the project; A.L. and A.B. designed the experiments; A.L. performed the majority of the experiments, with the help of A.B.; H.K. performed the computational analysis of RNA-seq, ChIP-seq experiments with inputs from A.L. and A.B.; C.M. performed smRNA-FISH; R.B. performed image analysis; A.R. performed the RACE-PCR experiments; H.J.W. performed 4C-seq analysis; S.M. performed the coding potential analysis; B.B. and P.G. performed and analyzed the MinION experiment; R.T. performed the computational analysis on the human gastrula data with the supervision of H.K.; K.M.P. conducted GATA4/6 ChIP-seq experiments; J.H. contributed to some aspects of the GATA4/6 ChIP-seq data and conservation analysis; R.M. gave critical input to the work and supervised some of the GATA4/6 ChIP-seq experiments; F.M gave critical input to the work and supervised the 4C data analysis; D.N. gave critical input to the work and supervised the coding potential analysis; D.M. performed the mass spectrometry analysis; J.R. gave critical input to the work and supervised the FISH experiments; A.L., A.B. and A.M. wrote the manuscript with critical inputs from all authors. A.M. supervised the project.

---

**Figure supplement 1:**
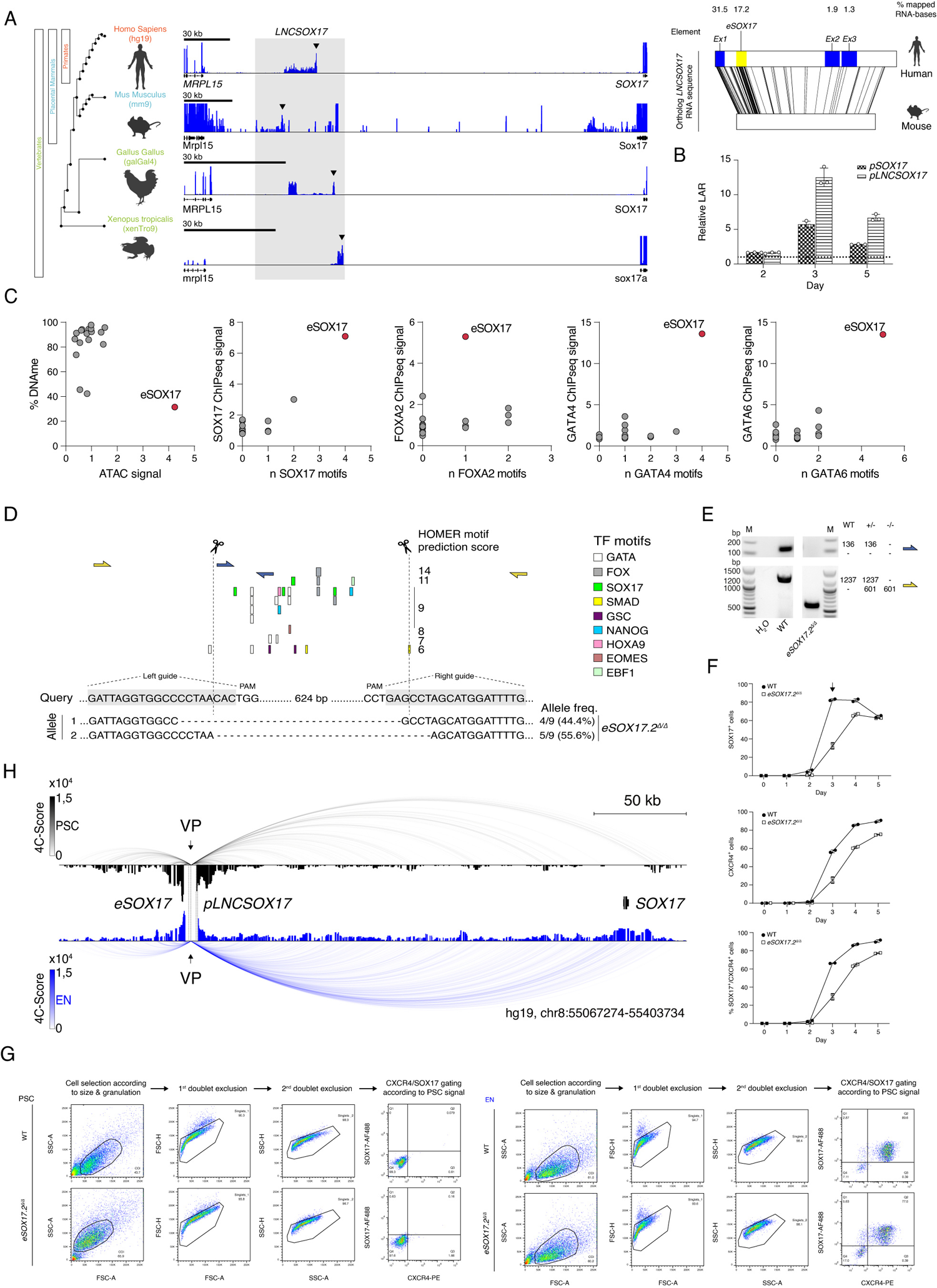
Functional characterization of the *SOX17* distal regulatory elements. **(A)** Left panel shows the identification of an unannotated transcript at the *SOX17* locus in embryonic stage RNAseq datasets across vertebrates (human (dEN), mouse (dEN), chicken (HH4) and frog (XT11)). Note how the relative position of the non-coding element (between *SOX17* and *MRPL15* genes) is conserved in all analyzed species. See Table 1 for the references to the datasets used in this analysis. The right panel indicates the EMBOSS Water alignment for the human vs. mouse *LNCSOX17* sequence. Numbers displayed are percentages per element of mapped bases including substitutions but excluding indels from the human sequence, where mapping is greater than or equal to 20 bases in length (black lines). **(B)** Firefly luciferase assay from either *pLNCSOX17* (hg19, chr8:55140804-55141678) or *pSOX17* (SOX17 promoter) (hg19, chr8:55365464-55370513) at day 2, 3 or 5 of EN differentiation. Values are calculated as luciferase activity ratio (LAR) between firefly and renilla signal, finally normalized on the empty vector background and day 0 baseline signal. Bars indicate mean values, error bars show standard deviation (SD) across three independent experiments. Individual data points are displayed. Raw measurements are reported in Table 1. **(C)** Scatter plots displaying DNA methylation levels, ATAC signal, endoderm TFs occupancy as measured by ChIP-seq and TFs binding motifs abundance at the *LNCSOX17* locus in EN cells. The *LNCSOX17* transcribed region was binned into 18 bins (dots) of the same size, including *eSOX17* (red dot). Note how *eSOX17* is depleted of DNA methylation and enriched in ATAC signal, endoderm TFs binding motifs and actual TFs occupancy as compared to the rest of the transcribed region, indicating a specific enhancer identity. **(D)** Schematic of the Cas9 based *eSOX17.2* gene ablation strategy. Genotyping PCR-products are depicted in Figure supplement 1E. sgRNA sequences are highlighted in grey while Cas9 targeting sites are depicted by dashed lines. Sanger sequencing results are summarized below the query-sequence and detected allele-frequency are displayed on the side for each respective genotype. **(E)** Genotyping PCR-products, generated by two different primer-pairs to profile *eSOX17.2* gene ablation (see schematic in Figure supplement 1D). Expected amplicon-sizes within a particular genetic background are shown on the side of the agarose gel-picture. **(F)** Line plots showing percentages of either SOX17^+^, CXCR4^+^ or SOX17^+^/CXCR4^+^ cell-populations at specified time-points during EN differentiation as measured by FACS in wild-type and eSOX17.2 knock-out cells. Symbols indicate the mean and error bars indicate SD across two independent experiments. Individual data points are displayed. **(G)** Gating strategy to quantify the percentages of SOX17 and CXCR4 positive populations used in this study in wild-type and corresponding gating strategy in the *eSOX17.2* knock-out. **(H)** 4Cseq of PSC (black) and EN (blue) at the *SOX17*-locus. Normalized interaction-scores displayed as arcs and histogram-profiles utilizing *eSOX17* as viewpoint (VP). *eSOX17* and *pLNCSOX17* are indicated by dotted lines.

**Figure supplement 2:**
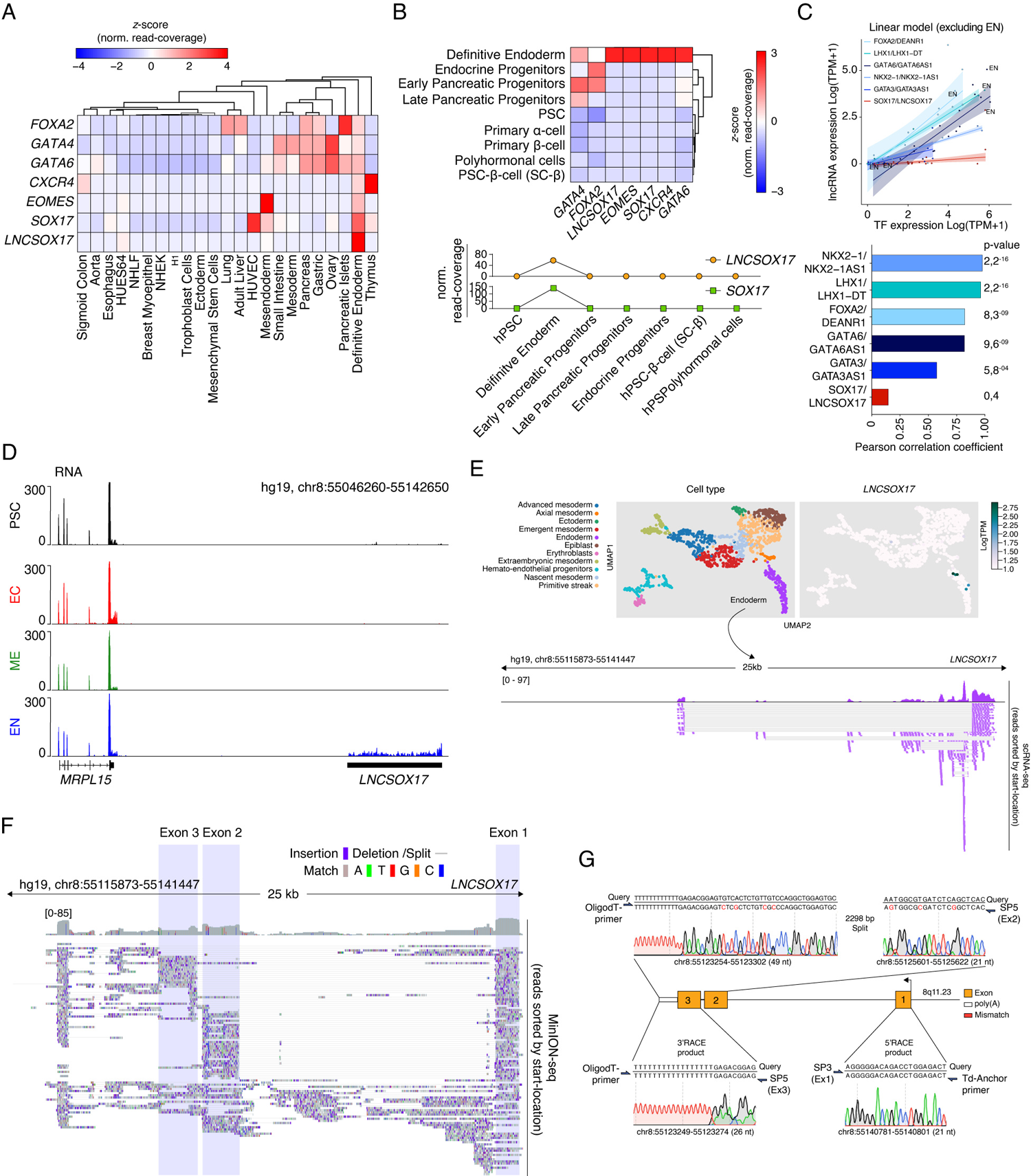
*LNCSOX17* tissue distribution and structural characterization. **(A)** Heatmap showing row-normalized *z*-scores of EN marker genes and *LNCSOX17* expression across various embryonic and adult tissues as measured by RNA-seq. Trees represent hierarchical clustering (based on Euclidean distance). RNA-seq expression values are calculated as normalized row-read-coverage. Note the high specificity of *LNCSOX17* expression as compared to a much broader expression of endodermal transcription factors, including *SOX17*. A list of the curated dataset from the Roadmap Epigenome project used in this heatmap is provided in Table 1. **(B)** Heatmap showing column-normalized *z*-scores of EN marker genes and *LNCSOX17* expression during *in vitro* pancreatic lineage differentiated cell types including primary isolated α- and β-cells as measured by RNA-seq (*80*) (upper panel). Trees represent hierarchical clustering (based on Euclidean distance). RNA-seq expression values are calculated as normalized row-read-coverage. *LNCSOX17* and *SOX17* expression profiles plotted as row-read-coverage normalized *z*-scores of the indicated *in vitro* generated cell type during pancreatic differentiation (lower panel). Note the transient and restricted expression of *LNCSOX17* specifically in definitive endoderm. **(C)** Scatter plot showing the expression of a set of endoderm lncRNAs (*DEANR1, LHX1-DT, GATA6AS1, NKX2-1AS1, GATA3AS1, LNCSOX17*) and the corresponding TFs (*FOXA2, LHX1, GATA6, NKX2-1, GATA3, SOX17*) in the same set of EN tissues of Figure 2B. A linear model excluding the expression in EN was fit for each lncRNA-TF couple. Pearson correlation coefficients as well as corresponding p-values are displayed in the bar-plot (bottom panel). Note that the *LNCSOX17-SOX17* couple has the lowest degree of tissue co-expression. **(D)** Genome browser tracks displaying RNA levels at *LNCSOX17* locus in PSCs and the three germ layers. Note *LNCSOX17* expression specificity as compared to *MRPL15*. **(E)** Uniform Manifold Approximation and Projection for Dimension Reduction (UMAPs) showing cell states (upper left panel) and *LNCSOX17* expression (upper right panel) in cells derived from a human gastrulating embryo (*44*). ScRNAseq track from cells belonging to the endoderm cluster showing reads mapping to the *LNCSOX17* locus (bottom panel). **(F)** MinIONseq reads track showing *LNCSOX17* coverage and structure in endodermal cells. Sequencing read distribution histogram (top) and individual reads sorted by their start-location (bottom) are displayed. Exon 1, 2 and 3 are highlighted by shading boxes. Sequence mismatches and matches are color coded as described. Split-reads and deletions are shown as thin horizontal lines. **(G)** Sanger sequencing of 3’/5’ RACE PCR products. Amplicon specific sequencing results are shown below the query sequence (hg19). Sequencing mismatches are highlighted in red. Primer pairs relative positions used for the PCRs are shown for each product. Sanger sequencing chromatogram color code is used to show the raw reads data.

**Figure supplement 3:**
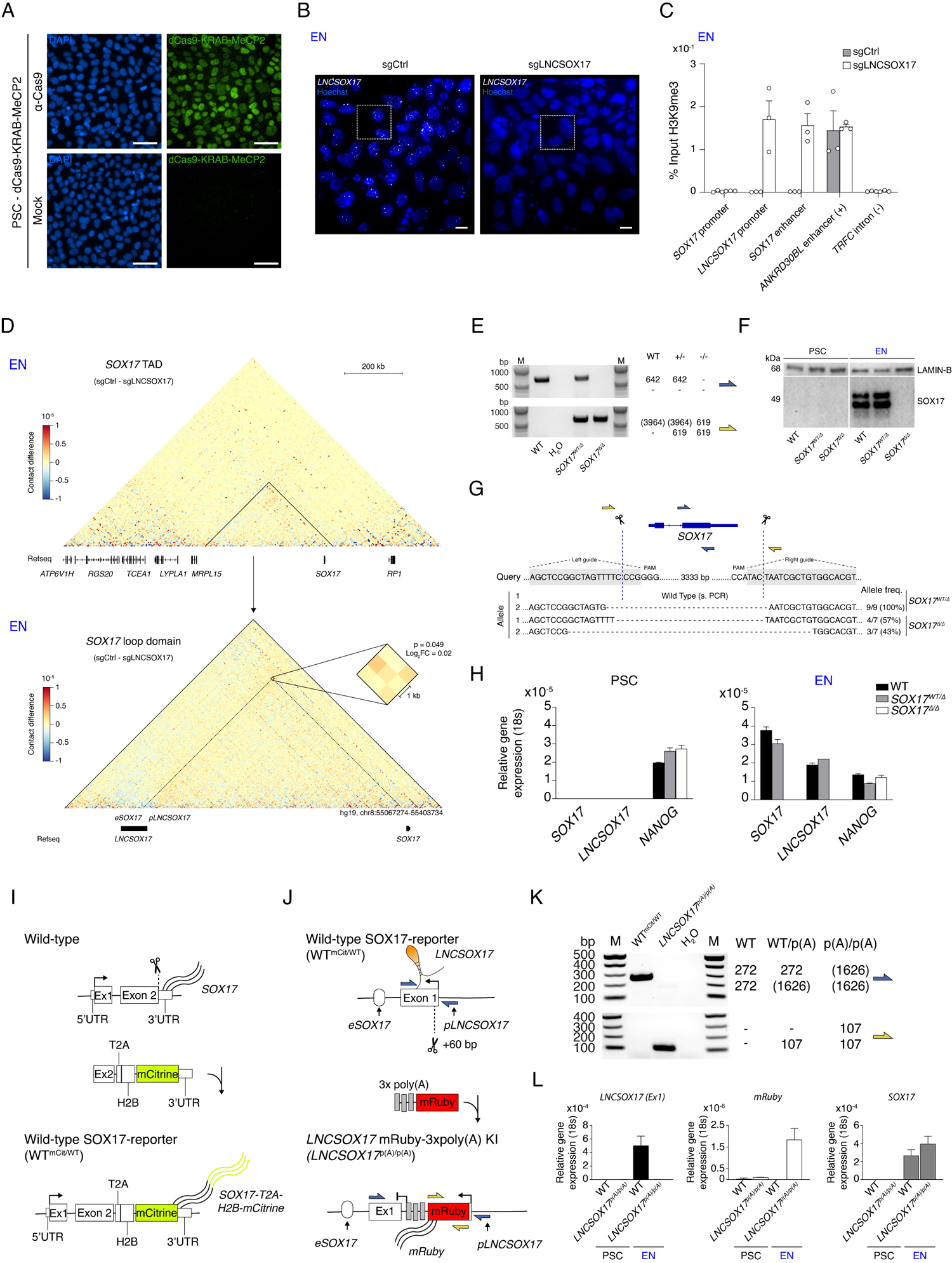
SOX17 and *LNCSOX17* reciprocal gene expression regulation. **(A)** IF-staining for dCas9 in PSCs expressing dCas9-KRAB-MeCP2 and counter-stained with DAPI. Mock samples represent secondary antibody only controls. Scale bars, 50 µm. **(B)** smRNA-FISH of LNCSOX17 in sgCtrl (left) and sgLNCSOX17 (right) EN cells counter-stained with Hoechst. Scale bars, 10 µm. Magnified regions are shown in Figure 3C. **(C)** H3K9me3 ChIP-qPCR enrichment percentages over input is represented at different regions of the genome in sgCtrl and sgLNCSOX17 endoderm cells. Bars indicate mean values, error bars indicate the SD across three independent experiments. See raw data in Table 4. **(D)** Capture Hi-C sequencing subtraction map of the EN sgCtrl-sgLNCSOX17 at the *SOX17* locus and at the SOX17 loop domain. *eSOX17* loop interaction with SOX17 promoter is shown in the magnification and highlighted by the dotted lines (significance threshold: log_2_FC ± 0.5, p < 0.01). **(E)** Genotyping PCR-products, generated by two different primer-pairs to profile *SOX17* gene ablation (see schematic in Figure supplement 3G). Expected amplicon-sizes within a particular genetic background are shown on the side of the agarose gel-picture. **(F)** Western Blot showing SOX17 levels in PSCs and EN cells for the three indicated genotypes. LAMIN-B is used as loading control. **(G)** Schematic of the Cas9 based *SOX17* gene ablation strategy. Genotyping PCR-products are depicted in Figure supplement 3E. sgRNA sequences are highlighted in grey while Cas9 targeting sites are depicted by dashed lines. Sanger sequencing results are summarized below the query-sequence and detected allele-frequency are displayed on the side for each respective genotype. **(H)** qRT-PCR showing *SOX17*, *LNCSOX17* and *NANOG* expression in PSCs and EN cells for the three indicated genotypes. Fold change is calculated relative to the *18s* housekeeping gene. Bar indicate the means, error bars represent SD across three independent replicates. **(I)** Schematic of the strategy to generate the SOX17-mCitrine reporter cell line. **(J)** Schematic of the strategy to generate the LNCSOX17^p(A)/p(A)^ cell line. **(K)** Genotyping PCR-products, generated by two different primer-pairs to profile the early poly(A) knock-in. Expected amplicon-sizes within a particular genetic background are shown on the side of the agarose gel-picture. **(L)** qRT-PCR showing *LNCSOX17, mRuby* and *SOX17* expression in PSCs and EN cells for the two indicated genotypes. Fold change is calculated relative to the *18s* housekeeping gene. Bar indicate the means, error bars represent SD across three independent replicates.

**Figure supplement 4:**
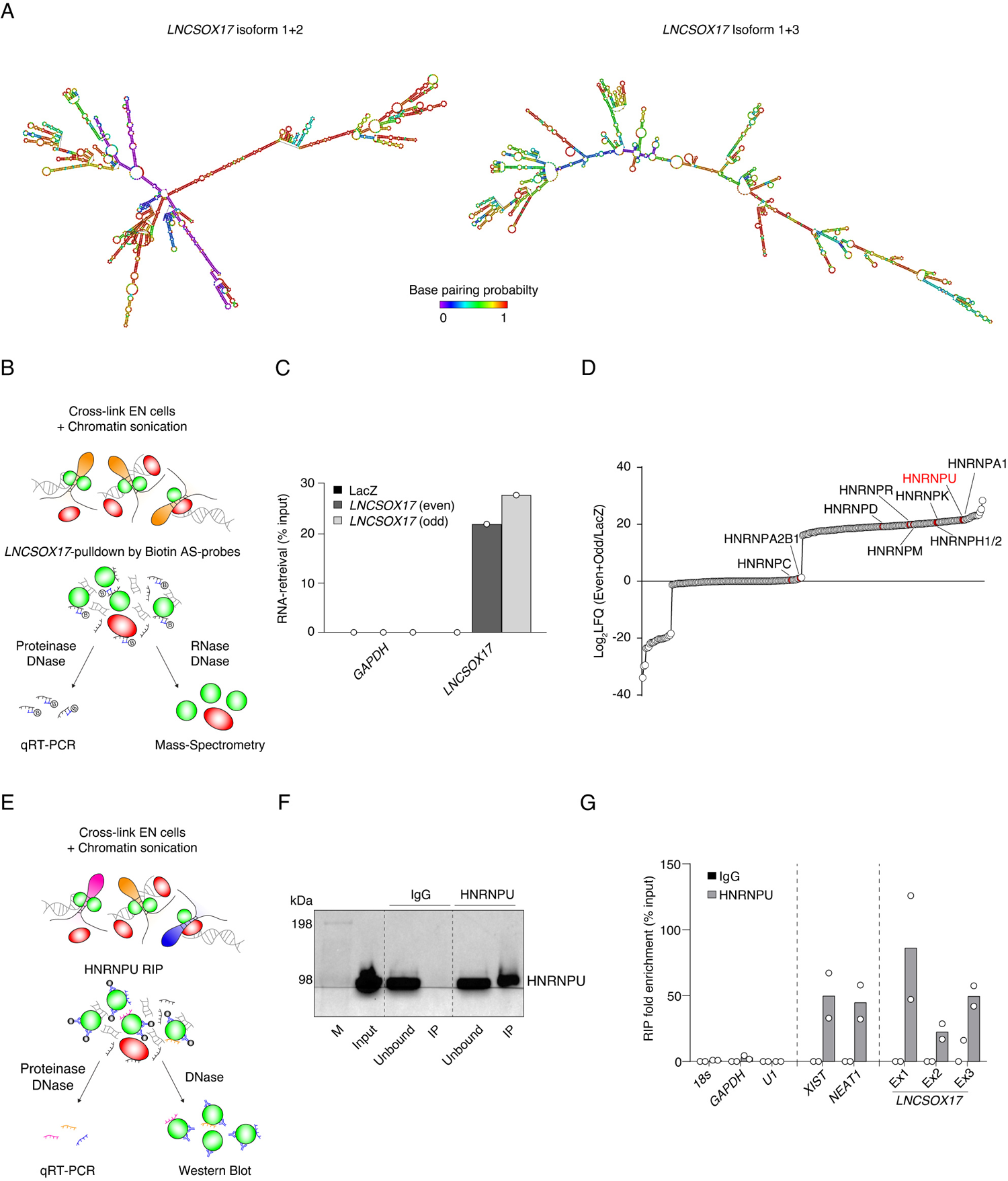
*LNCSOX17* interacts with HNRNPU. **(A)** RNA secondary structure predictions for *LNCSOX17* Ex1+2 and Ex1+3 isoforms as computed by the Vienna RNA Websuite tool. Base pairing probability is displayed with a color scale. **(B)** Schematic representation of *LNCSOX17* RNA-pulldown experimental workflow. **(C)** *LNCSOX17*-pulldown qPCR showing enrichment of *LNCSOX17* in both Even and Odd probe sets samples as compared to LacZ in endoderm cells. *GAPDH* RNA has been used as a negative control. **(D)** Ranked *LNCSOX17* protein interaction partners as measured by mass spectrometry. Log_2_LFQ (Even+Odd/LacZ) has been calculated for each measured peptide to highlight protein partners which are enriched in Even and Odd samples as compared to LacZ sample (positive values). LFQ, Label Free Quantification. **(E)** Schematic representation of HNRNPU RNA immunoprecipitation (RIP) experimental workflow. **(F)** Western Blot showing HNRNPU enrichment in the HNRNPU IP as compared to IgG control. Unbound sample represents post-IP supernatant. **(G)** RIP-qPCR showing enrichment of *LNCSOX17* in HNRNPU IP as compared to IgG in endoderm cells. *18S*, *GAPDH*, and *U1* RNAs have been used as negative controls, while *XIST* and *NEAT1* as positive controls. Exon-specific primer pairs have been used to probe *LNCSOX17*-HNRNPU interaction. Dots indicate two independent biological replicates.

**Figure supplement 5:**
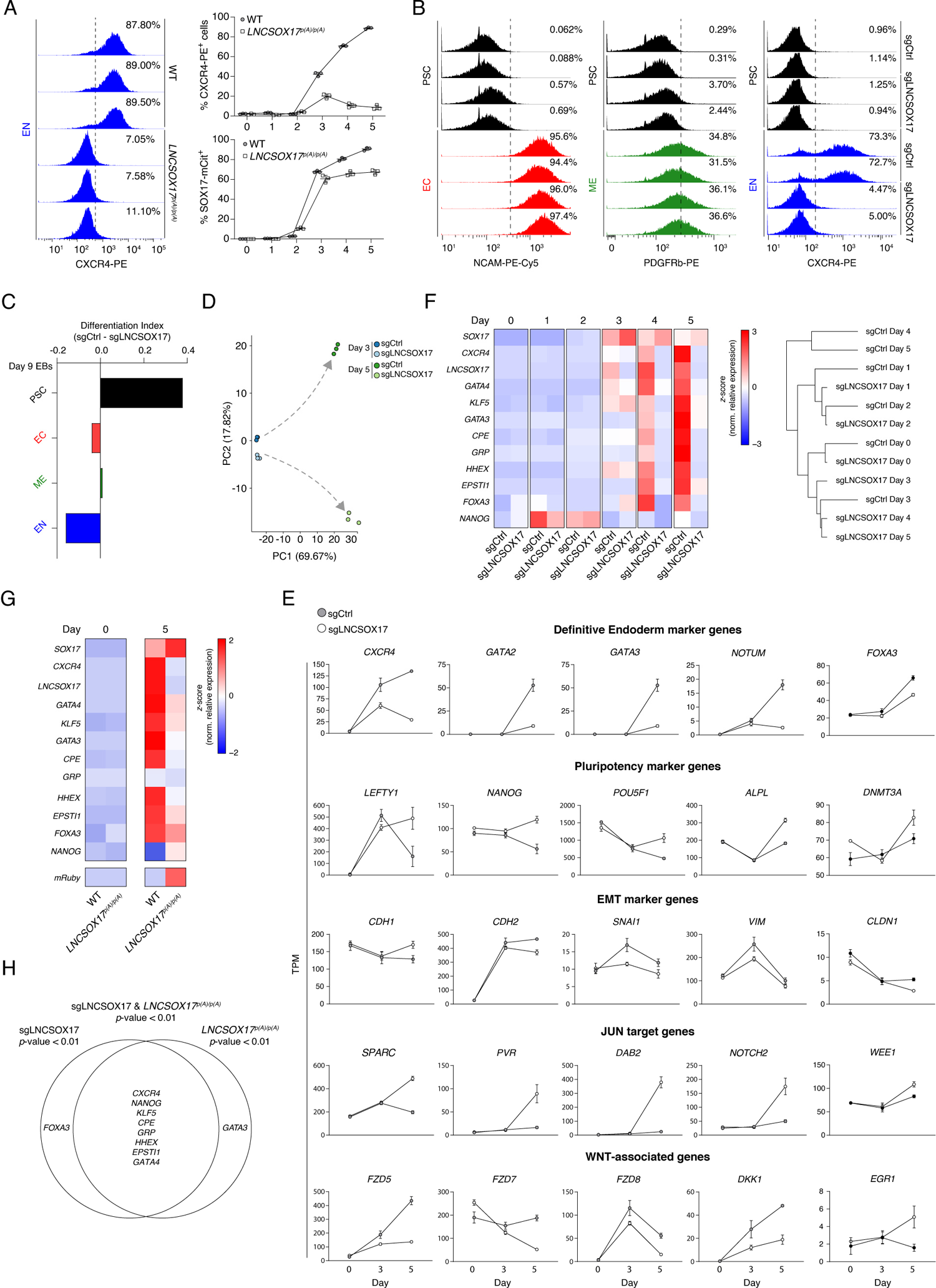
Molecular phenotypes associated with the loss of *LNCSOX17* **(A)** FACS histograms showing percentages of CXCR4^+^ cells during EN differentiation of wild-type and LNCSOX17^p(A)/p(A)^ cell lines (left), Sample sizes are normalized to 10000 cells /sample. In the right panel, FACS time-course experiment showing percentages of CXCR4^+^ and SOX17-mCitrine^+^ cells during EN differentiation of wild-type and LNCSOX17^p(A)/p(A)^ cell lines. Three independent replicates are displayed. **(B)** FACS histograms showing percentages of successfully differentiated cells during directed differentiation of sgCtrl and sgLNCSOX17 lines towards the three germ layers. Sample sizes are normalized to 8000 cells /sample. Two independent replicates are displayed. Note how no difference in percentages of differentiated cells is observed for the ectodermal and mesodermal trajectories, while a strong reduction is present in sgLNCSOX17 cells differentiating towards EN. **(C)** ScoreCard assay displaying differentiation index in sgCtrl or sgLNCSOX17 day 9 differentiated embryoid bodies (EBs) (*n*=48 EBs per line). **(D)** PCA of the 1000 most variable genes between day 3 and day 5 differentiation of sgCtrl and sgLNCSOX17 as measured by RNA-seq. Gray dashed arrows indicate the two divergent transcriptomic trajectories. **(E)** Line plots displaying time-resolved expression of selected marker genes in sgCtrl and sgLNCSOX17 cells as measured by RNA-seq (TPM). Marker genes categories are indicated. Symbols indicate mean values and error bars represent SD across three independent experiments. EMT, epithelial-to-mesenchymal transition. **(F)** Heatmap showing time-resolved qRT-PCR for endoderm specific marker genes during EN differentiation in sgCtrl and sgLNCSOX17 cell lines (left) and corresponding hierarchical clustering tree (Euclidean distance) (right). **(F)** Heatmap showing qRT-PCR for endoderm specific marker genes during EN differentiation in WT and LNCSOX17^p(A)/p(A)^ cell lines. **(G)** Venn diagram displaying the intersection between genes differentially expressed in the sgLNCSOX17 and LNCSOX17^p(A)/p(A)^ EN cells. Significantly downregulated (p<0,01) in both conditions are displayed in the intersection.

**Figure supplement 6:**
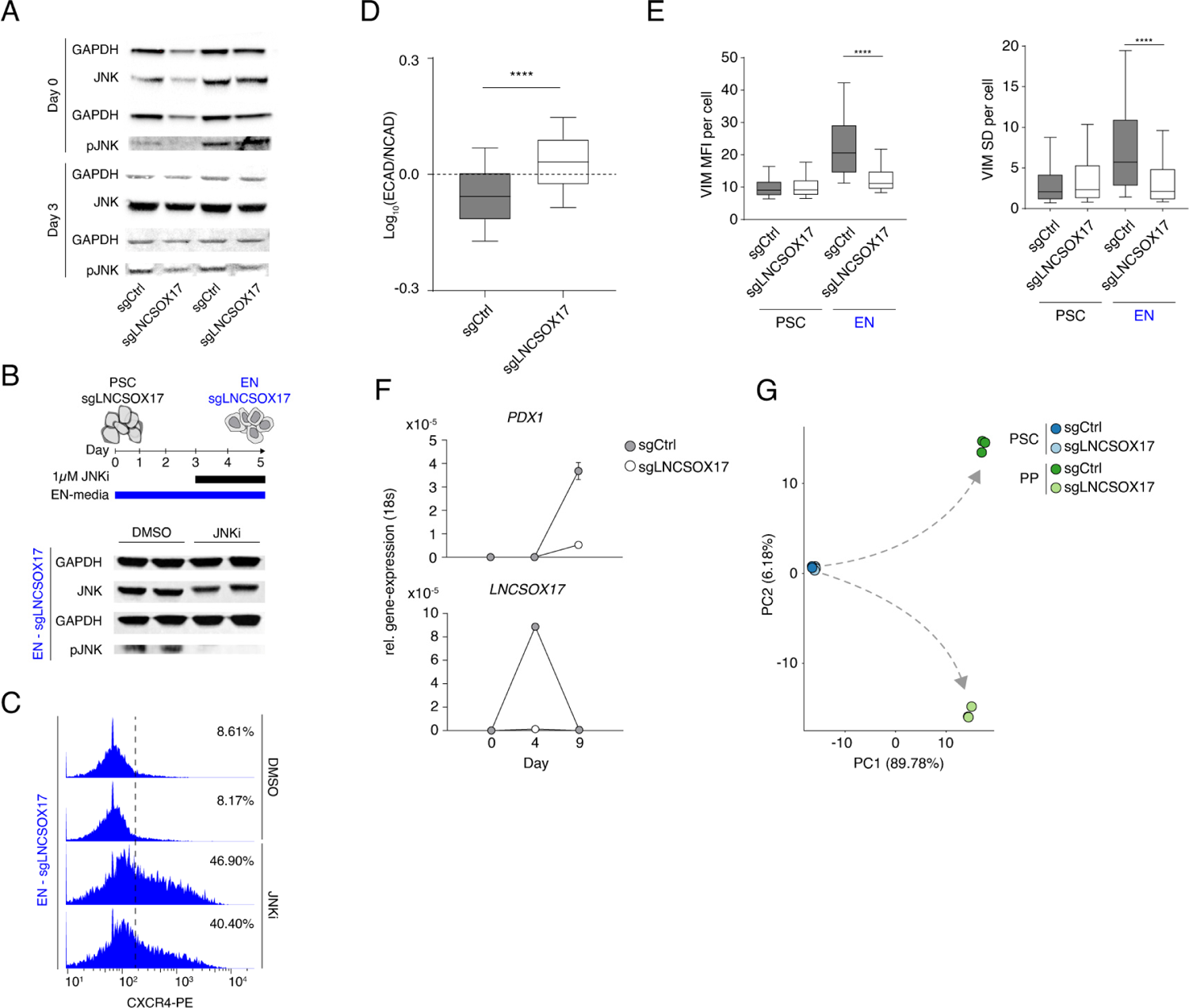
Cellular phenotypes associated with the loss of *LNCSOX17*. **(A)** Western blot showing the levels of JNK and pJNK during EN differentiation in sgCtrl and sgLNCSOX17 cell lines. Two independent replicates are displayed. GAPDH is used as loading control. **(B)** Western blot showing the levels of JNK and pJNK during EN differentiation in sgLNCSOX17 cell line in the presence or absence of JNK Inhibitor XVI from day 3 of EN differentiation (see schematic). Two independent replicates are displayed. GAPDH is used as loading control. **(C)** FACS histograms showing percentages of successfully differentiated cells during directed differentiation of sgLNCSOX17 line towards EN in the presence or absence of JNK Inhibitor XVI as measured by CXCR4^+^ cells, sample sizes are normalized to 8000 cells /sample. Two independent replicates are displayed. Note how the treatment with JNKi partially rescues the differentiation phenotype in cells lacking *LNCSOX17*. **(D)** IF stainings (Figure 4D) quantification showing Log_10_ (ECAD/NCAD) signal ratios from sgCtrl (*n*=28002) and sgLNCSOX17 (*n*=24284) EN cells. Boxes indicate 25^th^ and 75^th^ percentiles, central line indicate the median and whiskers show SD; p-value < 0,0001. List of values for each cell are shown in (Table 4). **(E)** IF staining (Figure 4E) quantification of per cell VIM signal (left) or VIM SD (right) of PSCs or EN sgCtrl (*n*PSCs=3058, *n*EN=11425) and sgLNCSOX17 (*n*PSCs=2329, *n*EN=14155) cells. VIM signal SD in the cellular compartment is used here as a measure of cellular polarization (s. methods). Boxes indicate 25^th^ and 75^th^ percentiles, central line indicate the median and whiskers show 10^th^ and 90^th^ percentiles; p-value < 0,0001 for both plots. List of values for each cell are shown in (Table 4). **(F)** qRT-PCR showing the expression of *PDX1* (upper panel) and *LNCSOX17* (lower panel) during PP differentiation of sgCtrl and sgLNCSOX17 cells. Fold change is calculated relative to the *18s* housekeeping gene. Symbols indicate mean values and error bars indicate SD across three independent experiments. **(G)** PCA of the 1000 most variable genes between sgCtrl and sgLNCSOX17 in PSCs and PP as measured by RNA-seq. Gray dashed arrows indicate the two divergent transcriptomic trajectories.

## Acknowledgements

We are grateful for the support and feedback received by all Meissner Lab members during the project development. Special recognition to T. Aktas and I.A. Ilik for experimental advising. B. Lukaszewska-McGreal for support with the mass spectrometry experiment. R.D. Acemel for suggestions regarding the virtual 4C analysis. I. Ulitsky for advice with the conservation analysis. B. Fauler for help with microscopy; D. Ibrahim for fruitful feedback; MPIMG Seq-Core for NGS support.

## Authors’ information

Landshammer Alexandro and Bolondi Adriano contributed equally to this work.

## TABLE DESCRIPTIONS

**Table 1:**

**-** List of external data sets used in this study.
**-** Luciferase assay raw and normalized data for eSOX17, eSOX17.1, eSOX17.1, pSOX17 and pLNCSOX17 of respective day of endodermal differentiation.
**-** MinION split reads extracted from IGV junctions-track BAM files.
**-** *LNCSOX17* isoform sequences.

**Table 2:**

- RNAseq data set with TPM values from undifferentiated (iPSC), day 3 and day 5 endoderm differentiated sgCtrl and sgLNCSOX17 CRISPRi cells.
**-** Differential gene expression analysis of RNAseq data between sgCtrl and sgLNCSOX17 from respective days of endoderm differentiated cells.
**-** RNAseq data sets with TPM values from day 9 pancreatic precursor (PP) differentiated sgCtrl and sgLNCSOX17 CRISPRi cells.
**-** Differential gene expression analysis of RNAseq data between sgCtrl and sgLNCSOX17 from respective days of pancreatic precursor differentiation.

**Table 3:**

- Taqman qRT-PCR probes used in this study.
**-** qRT-PCR primers, sgRNA oligonucleotides, cloning and genotyping primers used in this study.
**-** Antibodies and their respective application specific dilutions used in this study.
**-** *LNCSOX17* exonic smFISH probes and their sequence used in this study.
**-** *SOX17* locus capture HiC probes and their sequence used in this study.
**-** RNA-pulldown probes and their sequences used in this study.

**Table 4:**

- H3K9me3 ChIP-qPCR raw and normalized data from sgCtrl and sgLNCSOX17 day 5 endoderm cells.
**-** Raw data of Log_10_ (ECAD/NCAD) signal ratios and respective quantification from day 5 sgCtrl and sgLNCSOX17 endoderm cells.
**-** Raw data of VIMENTIN signal and respective quantification from sgCtrl and sgLNCSOX17 undifferentiated iPSCs and day 5 endoderm cells.
**-** Raw data of PDX1 signal and respective quantification from day 9 sgCtrl and sgLNCSOX17 pancreatic progenitor cells.

